# Translational modelling of low and medium intensity transcranial magnetic stimulation from rodents to humans

**DOI:** 10.1101/2024.04.27.591424

**Authors:** Samuel J. Bolland, Maxim Goryachev, Alexander Opitz, Alexander D Tang, Jennifer Rodger

**Affiliations:** Experimental and Regenerative Neurosciences, School of Biological Sciences, University of Western Australia, 35 Stirling Highway (M317), Crawley, 6009, WA, Australia; Perron Institute for Neurological and Translational Sciences, 8 Verdun Street, Nedlands, 6008, WA, Australia; Quantum Technologies and Dark Matter Laboratory, Department of Physics, University of Western Australia, 35 Stirling Highway, Crawley, Western Australia 6009, Australia; School of Biomedical Sciences, University of Western Australia, 35 Stirling Highway (M317), Crawley, 6009, WA, Australia; Department of Biomedical Engineering, University of Minnesota, Minneapolis, MN 55455, USA

**Keywords:** Finite element method, repetitive transcranial magnetic stimulation, translational model, electrical field distribution, magnetic field distribution, animal model, cross-species model

## Abstract

**Background:** Rodent models using subthreshold intensities of transcranial magnetic stimulation (TMS) have provided insight into the biological mechanisms of TMS but often differ from human studies in the intensity of the electric field (E-field) induced in the brain.

**Objective:** To develop a finite element method model as a guide for translation between low and medium intensity TMS rodent studies and high intensity TMS studies in humans.

**Methods:** FEM models using three head models (mouse, rat, and human), and eight TMS coils were developed to simulate the magnetic flux density (B-field) and E-field values induced by three intensities.

**Results:** In the mouse brain, maximum B-fields ranged from 0.00675 T to 0.936 T and maximum E-field of 0.231 V/m to 60.40 V/m E-field. In the rat brains maximum B-fields ranged from of 0.00696 T to 0.567 T and maximum E-fields of 0.144 V/m to 97.2 V/m. In the human brain, the S90 Standard coil could be used to induce a maximum B-field of 0.643 T and E-field of 241 V/m, while the MC-B70 coil induced 0.564 T B-field and 220 V/m E-field.

**Conclusions:** We have developed a novel FEM modelling tool that can help guide the replication of rodent studies using low intensity E-fields to human studies using commercial TMS coils. Modelling limitations include lack of data on dielectric values and CSF volumes for rodents and simplification of tissue geometry impacting E-field distribution, methods for mitigating these issues are discussed. A range of additional cross-species factors affecting the translation of E-fields were identified that will aid TMS E-field modelling in both humans and rodents. We present data that describes to what extent translation of brain region-specific E-field values from rodents to humans is possible and detail requirements for future improvement. A graphical abstract of the translational modelling pipeline from this study is provided below (Figure A.1).

**Highlights:**

- Clinical translation of rodent TMS studies is challenging due to the differences in coil and brain size and shape between rodents and humans.
- We have built a FEM model for the accurate replication of TMS-derived E-fields validated in rodent models in multiple brain regions in humans.
- This model is useful in designing stimulation parameters for humans based on rodent studies.
- This model is a critical part of a translational pipeline for evidence based TMS.

## 1: Introduction

Transcranial magnetic stimulation (TMS) is widely used to modulate neural networks by rapidly pulsing current through electromagnetic coils over the scalp to create rapidly changing magnetic flux density fields (B-fields) around the coils. These time varying B-fields induce electric fields (E-fields) within the electrically conductive brain, modulating cortical excitability (Fitzgerald, Fountain et al. 2006), neural plasticity (Goldsworthy, Hordacre et al. 2021), and connectivity (Beynel, Powers et al. 2020).

**Figure A.1:**
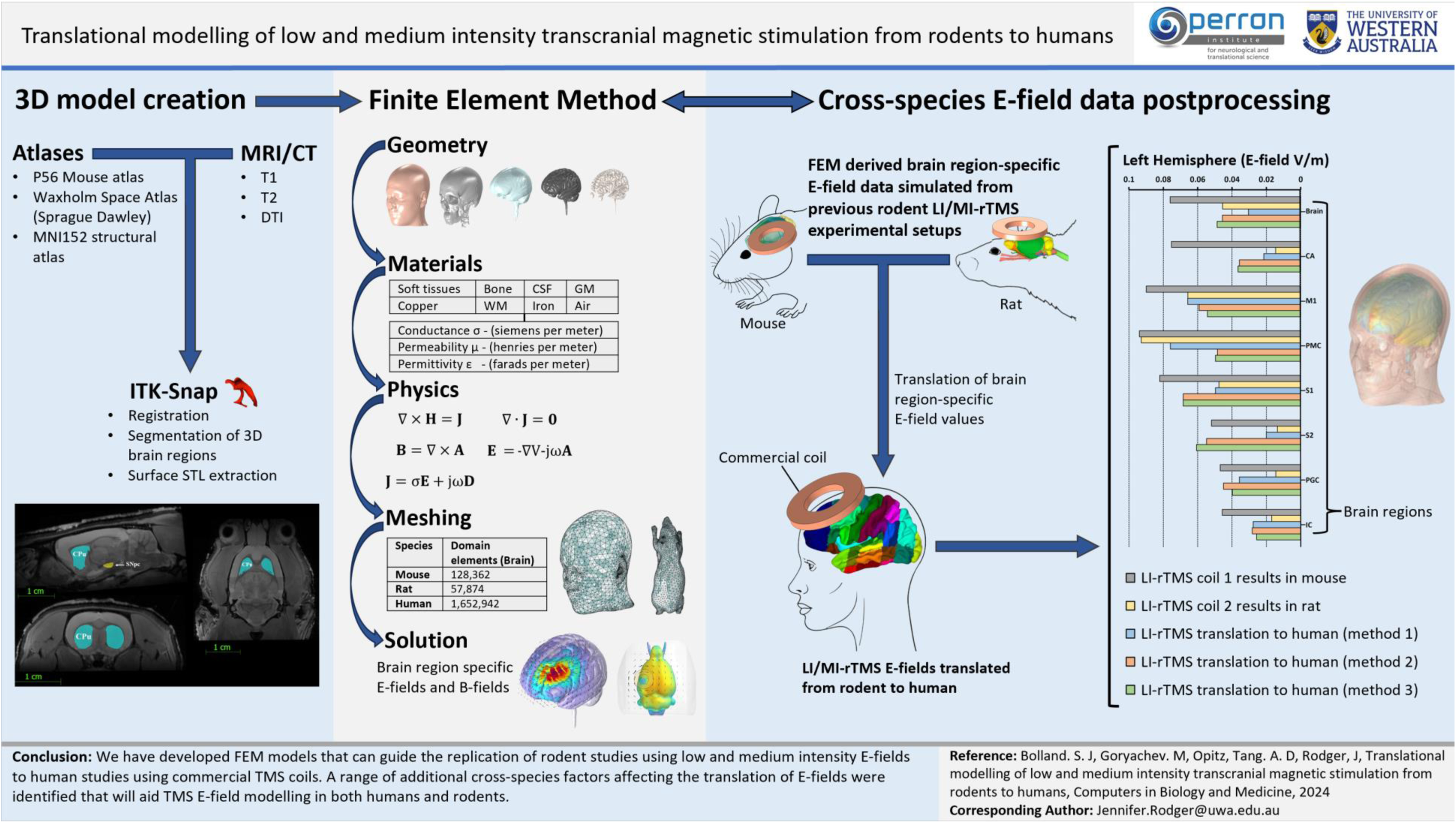
A translational modelling pipeline describing the use of structural imaging data for FEM modelling purposes and the comparison of brain region-specific rodent and human LI-rTMS E-field values.

To investigate biological mechanisms of TMS, an increasing number of studies take advantage of rodent models, in which invasive and post-mortem studies can be carried out (Tang, Thickbroom et al. 2017, Moretti and Rodger 2022). However, the small size of the rodent brain relative to the human brain has required compromise (Rodger and Sherrard 2015): either “high intensity” TMS (HI-TMS) is delivered using clinical equipment, resulting in a loss of focality (Salvador and Miranda 2009, Parthoens, Verhaeghe et al. 2016), or TMS is delivered using a miniaturised rodent-specific coil, resulting in a “low intensity” B-field and E-field (LI-TMS). HI-TMS in rodents has been defined as stimulation with a magnetic field of over 1 tesla produced at the base of the coil, whereas LI-TMS and medium intensity TMS (MI-TMS) produce a value between 0 and 0.05 tesla and 0.05 and 1 tesla respectively (Rodger and Sherrard 2015, Moretti and Rodger 2022). LI-TMS and MI-TMS are believed to induce subthreshold E-fields relative to the action potential threshold (i.e. not directly inducing neuronal spiking, but see Boyer, Baudin et al. (2022)). Nevertheless, LI-TMS and MI-TMS have been shown to induce synaptic mechanisms of plasticity (Tang, Bennett et al. 2021) and a range of biological effects have been observed including, but not limited to, structural reorganization (Makowiecki, Harvey et al. 2014), increased neurogenesis (Heath, Lindberg et al. 2018), gene regulation (Grehl, Viola et al. 2015), increased intracellular calcium (Grehl, Viola et al. 2015, Clarke, Penrose et al. 2017), altered neuron excitability (Tang, Lowe et al. 2016, Makowiecki, Garrett et al. 2018), and glial plasticity (Clarke, Penrose et al. 2017, Cullen, Senesi et al. 2019). These results suggest that TMS delivered at low intensity may activate mechanisms that are distinct from those involved in conventional high intensity rTMS, and may provide unique therapeutic avenues for specific disorders (Moretti and Rodger, 2022). An example of this is the specific impact of LI-rTMS on oligodendrocyte survival, which has led to a clinical trial for the treatment of multiple sclerosis (Cullen et al., 2019; Makowiecki et al., 2022). However, the differences in brain size and B-field intensity between humans and rodents in the literature and the complex relationship between E-field and biological tissue make translation of preclinical findings difficult. As a result, computer models simulating the physical phenomena behind TMS-brain interactions have emerged as useful tools to describe TMS induced electromagnetic field distribution in humans (Gomez-Tames, Laakso et al. 2020), rodent models (Tang, Bennett et al. 2021), and clinical TMS comparatively across different species (Alekseichuk, Mantell et al. 2019). Finite element method (FEM) modelling is an accurate way to simulate electromagnetic field distributions induced by TMS in the whole brain and within individual brain regions (Gomez-Tames, Laakso et al. 2020). However, there are limited attempts to describe the induced electromagnetic fields in the rodent head using the miniaturised LI-TMS and MI-TMS coils (Tang, Lowe et al. 2016, Madore, Poh et al. 2021, Moretti, Poh et al. 2021, Tang, Bennett et al. 2021, Zimdahl, Thomas et al. 2021). Development of a translational pipeline is necessary for prediction of TMS intensities and coil positioning to achieve homologous stimulation in brain regions of different species. No attempt to date has been made to directly compare commercially available human TMS coils with miniaturised rodent TMS coils.

To accurately represent the E-field in this FEM model, and as a form of future proofing the implementation of multiscale models, Faraday’s Law, Maxwell’s equations, and the constituent equations will be modified as little as possible within the COMSOL environment. Often, forms of quasistatic approximation (QSA) are employed for TMS stimulation due, in part, to an assumption that the ratio between head model sizes and electromagnetic wavelength (electrical size) leads to conductance being the dominant E-field determining dielectric value (AC/DC Module User’s Guide, pp. 113, COMSOL Multiphysics v.6.2.2), while permeability and permittivity are described as having relatively negligible impacts (Greiner 2012, Wang, Peterchev et al. 2024). QSA is a practical application for certain investigations, such as parameter sweeps where computational efficiency is key (Wang, Peterchev et al. 2024). However, use of QSA could neglect inductive, capacitive and propagation effects (Cvetković, Poljak et al. 2015) and the capacitive effect itself can create a difference of neuron level sense electric fields up to 22% (Gaugain, Quéguiner et al. 2023). Furthermore, the impact of permittivity and permeability in relation to electrical size is increased due to complex, non-planar, head geometry. However, our translational model avoids use of QSA to ensure that the E-field, B-field, H-field (magnetic field intensity), and D-field (electric displacement field) along with all three dielectric properties mentioned are mutually dependant to better inform E-field for a range of parameters and geometric scales. Supplementary Section S.1.2 detail the equations used in COMSOL to generate translational modelling solutions.

A range of cross-species differences make replicating rodent TMS studies in humans complex. To investigate to what degree this replication is possible we developed a modelling pipeline to translate rodent TMS studies to human clinical studies. This was achieved using FEM models to approximate brain region-specific electromagnetic field distributions from previous TMS rodent studies and from clinical human TMS settings. This translational model simulated an existing commercial coil and adjusted the stimulation parameters to reproduce LI-TMS and MI-TMS induced E-field distributions from rodent models in a human brain. E-field distributions will be described by maximum, minimum and average values in the whole brain and 26 comparative brain regions of interest. In doing this, we aim to provide a translational modelling pipeline, which can be adapted to simulate various experimental and clinical setups. We also aim to identify TMS modelling artifacts and cross-species translational issues and providing forms of mitigation. The purpose of this work is to aid in the acceleration of novel TMS treatments, which hold the potential to alleviate the suffering associated with neuropsychiatric disorders. Furthermore, providing an guide to better predicting E-field distribution helps further classify forms of neurostimulation by the magnitude and type of E-field induced, which could improve accuracy of future treatments as the E-field is understood to most directly mediate the effects of neurostimulation (Peterchev, Wagner et al. 2012).

In our models, stimulation is targeted to be unilateral to the left hemisphere, targeting the left brain hemisphere in the rodent models and the left dorsolateral prefrontal cortex (DLPFC) in humans due to its clinical preference for the treatment of depression (George, Nahas et al. 2000, Fitzgerald, Fountain et al. 2006, Fitzgerald, Hoy et al. 2009, Rusjan, Barr et al. 2010).

## 2: Materials and Methods

### 2.1 FEM modelling

FEM modelling was conducted in the frequency domain using COMSOL Multiphysics 6.1 (Burlington, NJ, USA), within the AC/DC Module’s Magnetic and Electric Fields physics interface, which solves Maxwell’s and the constitutive equations, to simulate the TMS induced B-field and E-field distributions in 3D mouse, rat, and human head models, with coil parameters based on previous relevant TMS research. The mouse, and human head models used in this paper are representative of widely used experimental models; the rat head model was custom built from MRI images. All coils were positioned unilaterally over the left side of the brain (over the DLPFC in the human model). Clean E-field distributions of all coils over a sphere can be seen in the Supplementary (Section S.1).

Modelling outputs were calculated using COMSOL’s magnetic and electric fields physics interface; Ampère’s Law and Current Conservation was used with the surrounding boundary being a cube consisting of surfaces with perfect magnetic conductance. Meshing was conducted with “extremely fine” as the element size, while the coil and core were meshed using the swept function. Only the biological tissues in the model have frequency dependent dielectric properties that alter the resulting E-field induced in the brain (Foster and Schwan 1986, Gabriel, Lau et al. 1996). Dielectric values for tissues in this study were obtained from the Foundation for Research on Information Technologies in Society dielectric tissue properties database (Hasgall, Di Gennaro et al. 2015).

### 2.2 Coil parameters

The rise time was calculated as the time across the section of the pulse where the voltage most rapidly changed between 0 volts and the peak voltage, or when the E-field would be at its greatest magnitude. TMS pulses in this study are translated from transient rise time values into the less computationally demanding frequency domain for modelling purposes (see section S.2), which is representative of a TMS pulse (Davey, Epstein et al. 2003, Davey and Riehl 2006, Deng, Lisanby et al. 2014). The term frequency is applied in this study to the period of stimulation in the frequency domain, generally frequency is used in clinical TMS applications to refer to the pattern used to deliver repetitive pulses. All commercial coil models were stimulated by outputs expected by The Magstim 200^2^ Stimulator (MagStim Company, Whitland, UK) and MagPro R30/X100 for Magventure (MagVenture A/S, Denmark), which have maximum stimulator outputs (100% MSO) of 165 A/µs^-1^ and 150 A/ µs^-1^ respectively. An Effective Nonlinear Magnetic Curves Calculator was used to produce a more realistic frequency domain response in iron-core coils (see Supplementary Materials, section S.3.1). An investigation into variances observed between iron-core and air-core MI-TMS rodent coils was conducted in COMSOL to validate modelling and laboratory results (see S.3.2). COMSOL’s Electric Circuit physics interface was used to produce coil excitation (see S.4). All coil parameters are described in Table 2.1 below.

**Table 2.1:**
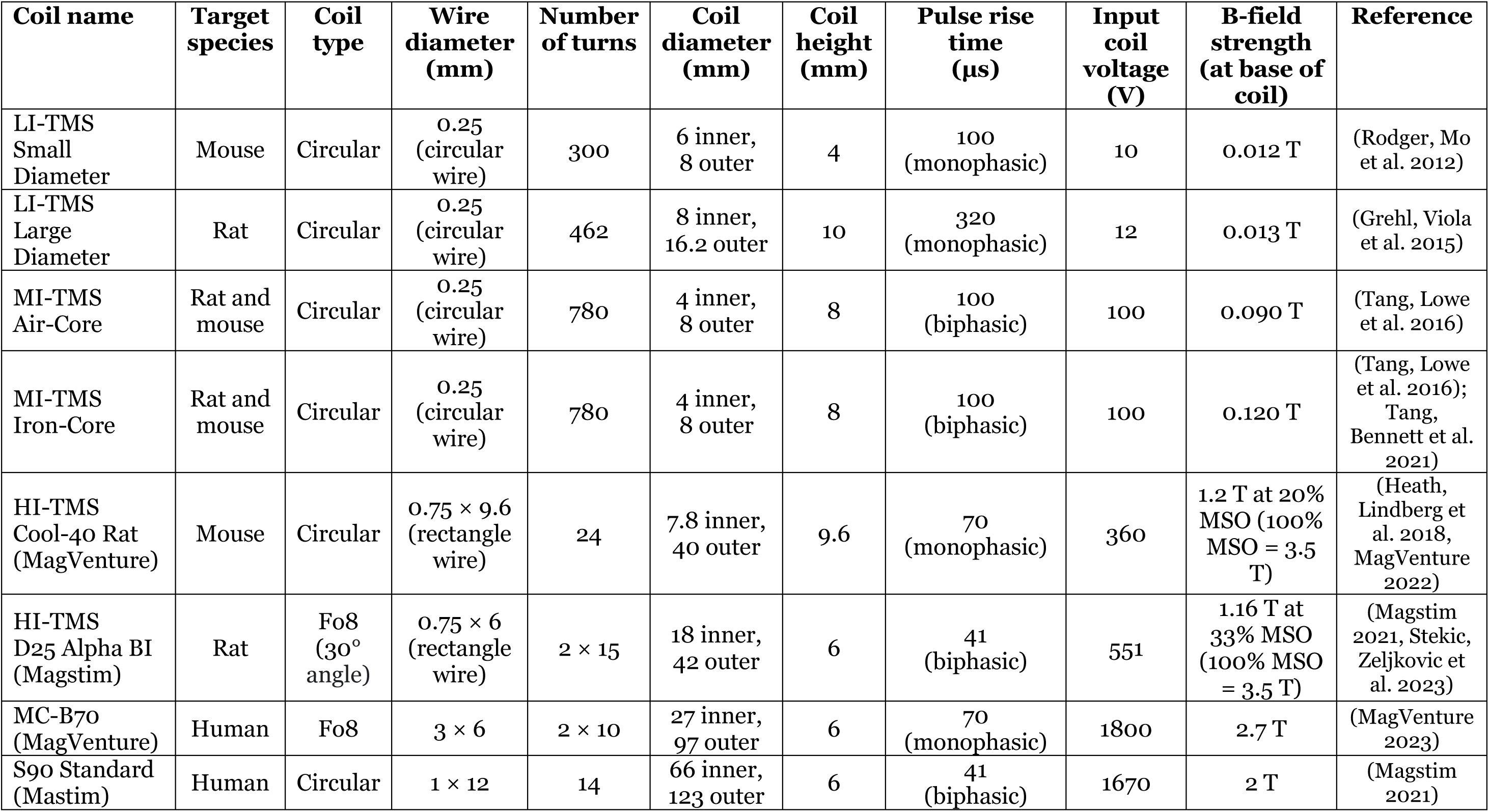
Coil parameters for each of the seven coils modelled in this study.

| Coil name | Target species | Coil type | Wire diameter (mm) | Number of turns | Coil diameter (mm) | Coil height (mm) | Pulse rise time (µs) | Input coil voltage (V) | B-field strength (at base of coil) | Reference |
| --- | --- | --- | --- | --- | --- | --- | --- | --- | --- | --- |
| LI-TMS Small Diameter | Mouse | Circular | 0.25 (circular wire) | 300 | 6 inner, 8 outer | 4 | 100 (monophasic) | 10 | 0.012 T | (Rodger, Mo et al. 2012) |
| LI-TMS Large Diameter | Rat | Circular | 0.25 (circular wire) | 462 | 8 inner, 16.2 outer | 10 | 320 (monophasic) | 12 | 0.013 T | (Grehl, Viola et al. 2015) |
| MI-TMS Air-Core | Rat and mouse | Circular | 0.25 (circular wire) | 780 | 4 inner, 8 outer | 8 | 100 (biphasic) | 100 | 0.090 T | (Tang, Lowe et al. 2016) |
| MI-TMS Iron-Core | Rat and mouse | Circular | 0.25 (circular wire) | 780 | 4 inner, 8 outer | 8 | 100 (biphasic) | 100 | 0.120 T | (Tang, Lowe et al. 2016); Tang, Bennett et al. 2021) |
| HI-TMS Cool-40 Rat (MagVenture) | Mouse | Circular | 0.75 × 9.6 (rectangle wire) | 24 | 7.8 inner, 40 outer | 9.6 | 70 (monophasic) | 360 | 1.2 T at 20% MSO (100% MSO = 3.5 T) | (Heath, Lindberg et al. 2018, MagVenture 2022) |
| HI-TMS D25 Alpha BI (Magstim) | Rat | Fo8 (30° angle) | 0.75 × 6 (rectangle wire) | 2 × 15 | 18 inner, 42 outer | 6 | 41 (biphasic) | 551 | 1.16 T at 33% MSO (100% MSO = 3.5 T) | (Magstim 2021, Stekic, Zeljkovic et al. 2023) |
| MC-B70 (MagVenture) | Human | Fo8 | 3 × 6 | 2 × 10 | 27 inner, 97 outer | 6 | 70 (monophasic) | 1800 | 2.7 T | (MagVenture 2023) |
| S90 Standard (Mastim) | Human | Circular | 1 × 12 | 14 | 66 inner, 123 outer | 6 | 41 (biphasic) | 1670 | 2 T | (Magstim 2021) |

Six rodent coil models were created to simulate TMS on either the mouse or rat head model depending on their use in previous studies. The rodent coils included in this study were, the 0.012 T LI-TMS Small Diameter Coil used previously on mice by Rodger, Mo et al. (2012), Makowiecki, Harvey et al. (2014), Yates, Martin-Iverson et al. (2014), and Poh, Harvey et al. (2018), the 0.013 T LI-TMS Large Diameter Coil used previously on rats by Grehl, Viola et al. (2015) and Seewoo, Feindel et al. (2018), the 0.090 T MI-TMS Air-Core Coil used previously on rats by (Tang, Lowe et al. 2016) and in mice by Heath, Lindberg et al. (2018), the 0.120 T MI-TMS Iron-Core Coil used previously on rats by Tang, Lowe et al. (2016) and on mice by Tang, Bennett et al. (2021), the HI-TMS Cool-40 Rat Coil (MagVenture A/S, Denmark) used previously on mice by Heath, Lindberg et al. (2018), and the HI-TMS D25 Alpha BI Coil (MagStim Company, Whitland, UK) used previously on rats by Stekic, Zeljkovic et al. (2023).

Two commercial human coil models were created to simulate TMS on the human head model. The human coils included in this study were the MC-B70 Coil (MagVenture A/S, Denmark) and the S90 Standard Coil (MagStim Company, Whitland, UK). The S90 Standard coil was selected to translate rodent LI-TMS and MI-TMS to a human model based on its circular shape and similar coil-to-brain ratio as the rodent coils. Details on all model coil placements relative to head models is provided in the Supplementary Material (section S.1.1).

Due to lack of publicly available rodent tissue-specific dielectric values, default isotropic conductivities were set for both rodents and human models (see Table 2.2) based on the IT’IS database (Hasgall, Di Gennaro et al. 2018), which is a replication of the efforts by the Gabriel & Gabriel database (Gabriel 1996). These values were previously used in human (Wagner, Zahn et al. 2004, Alekseichuk, Mantell et al. 2019) and non-human models (Nowak, Mix et al. 2011, Crowther, Hadimani et al. 2014, Alekseichuk, Mantell et al. 2019, Choung, Bhattacharjee et al. 2023, Jiang, Isenhart et al. 2023), as well as in the popular TMS simulation program SimNIBS (Saturnino, Puonti et al. 2019).

**Table 2.2:** Dielectric values for materials used in the construction of FEM models in this current study for all species.

| **Material type** | **Conductance**<br>($\sigma$ in siemens per meter) | **Permeability**<br>($\mu$ in henries per meter) | **Permittivity**<br>($\epsilon$ in farads per meter) |
| --- | --- | --- | --- |
| Soft tissues | 0.050 | 1 | $7.99 \times 10^2$ |
| Bone (skull) | 0.1 | 1 | $1.64 \times 10^2$ |
| Cerebrospinal fluid | 2.000 | 1 | $1.09 \times 10^2$ |
| Grey matter | 0.189 | 1 | $6.05 \times 10^2$ |
| White matter | 0.116 | 1 | $3.08 \times 10^2$ |
| Copper | $5.998 \times 10^7$ | 1 | 1 |
| Iron | $1.040 \times 10^7$ | $4.0 \times 10^3$ | 1 |
| Air | 0 | 1 | 1 |

### 2.3 Head models

3D FEM head models were constructed for a mouse, rat, and human using the CT scans of “Digimouse” (Dogdas, Stout et al. 2007) and the P56 mouse brain atlas (Lein, Hawrylycz et al. 2007) for the mouse, a T2 weighted anatomical MRI image for the rat and the CerebrA (MNI152) coordinate system for the human (Fonov, Evans et al. 2009, Manera, Dadar et al. 2020). Details on the production of these models can be found in the Supplementary Materials (section S.8).

Brain and head dimensions for each species are listed in Table 2.3 along with model mesh resolution. All FEM head models were meshed with triangle surface elements and tetrahedral volume elements.

**Table 2.3:**
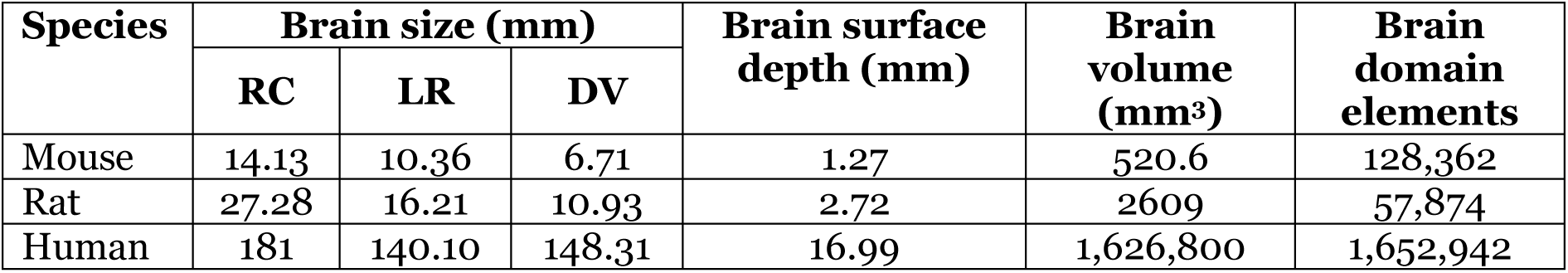
Dimensions for head and brain models used in this study (not including brain stem or spinal cord), including cortex surface depth from scalp surface (Brain surface depth) and mesh resolutions domain elements (tetrahedra).

### 2.4 Brain regions

Brain regions were sectioned out from the mouse, rat, and human head models and E-field values were obtained at 3D points 0.25 mm apart for the mouse and rat and 4 mm apart for the human (see Table 2.4).

**Table 2.4:** Comparative brain regions segmented out for the mouse, rat, and human 3D models. Comparative regions are aligned in rows and brain regions in head models were separated into left and right hemispheres where applicable.

| Mouse brain regions | Rat brain regions | Human brain regions |
| --- | --- | --- |
| CA (Hippocampal Region) | CA (Hippocampal Region) | CA (Hippocampal region) |
| MOp (Primary motor area) | MOp (Primary motor area) | M1 (Motor cortex) |
| MOs (Secondary motor cortex) | MOs (Secondary motor cortex) | PMC (Premotor cortex) |
| S1BF (Primary somatosensory cortex, barrel field) | S1BF (Primary somatosensory cortex, barrel field) | S1 (Somatosensory cortex)<sup>1</sup> |
| SSS (Supplementary somatosensory cortex) | S2 (Secondary somatosensory cortex) next to above in intensity | S2 (Secondary somatosensory cortex) |
| GU (Gustatory areas) | GI (Granular insular cortex) | PGC (Primary gustatory cortex) |
| VISC (Visceral area) | DI (Dysgranular insular cortex) | IC (Insular cortex) |
| AUDd (Dorsal auditory area) | AuD (Secondary auditory area, dorsal part) | A3 (Tertiary auditory cortex, parabelt) |
| AUDp (Primary auditory area) | Au1 (Primary auditory area) | A1 (Primary auditory cortex, core) |
| AUDv (Ventral auditory area) | AuV (Secondary auditory area, ventral part) | A2 (Secondary auditory cortex, belt) |
| VISal (Anterolateral visual area) | V2L (Secondary visual cortex, lateral area) | V3 (Tertiary visual cortex) |
| VISam (Anteromedial visual area) | V2M (Secondary visual cortex, medial area) | V2 (Secondary visual cortex) |
| VISp (Primary visual area) | V1 (Primary visual cortex) | V1 (Primary visual cortex) |
| ACAD (Anterior cingulate area, dorsal Part) | Cg1 (Cingulate area 1) | DLPFC (Dorsolateral prefrontal cortex) |
| ACAV (Anterior cingulate area, ventral Part) | Cg2 (Cingulate area 2) | vACC (Ventral anterior cingulate cortex) |
| PL (Prelimbic area) | PL (Prelimbic area) | VLPFC (Ventrolateral prefrontal cortex) |
| ORBL (Orbital area, lateral part) | Lateral orbital area (LO) | OPFc (Orbitofrontal prefrontal cortex) |
| ORBM (Orbital area, medial part) | Medial orbital area (MO) | dACC (dorsal anterior cingulate cortex) |
| ORBVL (Orbital area, ventrolateral part) | Ventrolateral orbital area (VLO) | POC (Primary olfactory cortex) |
| AID (Agranular insular area, dorsal part) | AID (Agranular insular area, dorsal part) | GIA (Granular insula area) |
| AIP (Agranular insular area, posterior part) | AIP (Agranular insular area, posterior part) | DIA (Dysgranular insula area) |
| AIV (Agranular insular area, ventral part) | AIV (Agranular insular area, ventral part) | AIA (Agranular insula area) |
| PTLP (Posterior parietal association areas, ventral part) | PPC (Parietal association cortex, posterior area) | PPC (Posterior parietal cortex) |
| TEA (Temporal association areas, ventral part) | TEA (Temporal association area) | TeA (Temporal association cortex) |
| PERI (Perirhinal area, ventral part) | PRC (Perirhinal cortex, areas 35 and 36) | PRC (Perirhinal cortex) |
| ECT (Entorhinal area) | ECT (Entorhinal area, medial and lateral) | EC (Entorhinal cortex) |

## 3: Results

Results of the LI-TMS Small Diameter, MI-TMS Air-Core, and HI-TMS Cool-Rat 40 coil simulations on the mouse head model are reported in Section 3.1. Section 3.2 reports the LI-TMS Large Diameter, MI-TMS Iron-Core, and HI-TMS D25 Alpha BI coil results simulated on the rat head model. The human coils, S90 and MC-B70, are reported in Section 3.3 at 50% MSO and Section 3.4 reports results from the S90 Standard coil, used to translate the electromagnetic field distributions from the two LI-TMS and three MI-TMS rodent coils. Results from SimNIBS (Version 4.0.0) for all the MC-B70 and S90 simulations are also reported in the Supplementary materials (section S.6 and Figure S.5) in support of the COMSOL models. Additionally, Section 3.5 describes a 10 mm section of brain surface parallel to the base of the coil for both the human and rat models to investigate the impact of human brain folding on E-field distribution. Peak value heat maps for rodent models are reported in Supplementary Figure S5.2. Mouse models used a range of 481,537 to 488,899 elements and 3,708,258 to 3,711,320 degrees of freedom. The rat models used a range of 547,542 to 658,001 elements and 4,221,065 to 4,983,552 degrees of freedom. Human models used a range of 3,589,778 to 3,601,241 elements and 27,547,009 to 27,633,685 degrees of freedom.

### 3.1 B-field and E-field results for rodent coils simulated on the mouse head model

The maximum induced B-field and E-field within the mouse brain was 6.75×10^-3^ T and 0.231 V/m respectively for the LI-TMS Small Diameter coil, 3.27×10^-2^ T and 1.820 V/m respectively for the MI-TMS Air-Core, 3.13×10^-2^ T and 1.456 V/m respectively for the MI-TMS Iron-Core, and 9.36×10^-1^ T and 60.40 V/m for the HI-TMS Cool-40 Rat coil (See Figure 3.3 for the MI-TMS Iron-core coil results and Figure 3.1 for all other mouse coil results). There was large peak in E-field values from all mouse coils found at lambda where the most caudal location along midline of the cortex meets the most rostral part of the midbrain forming an angled concave surface (Figure 3.2.), due to the tendency for electrical charge to gather at sharp angles between dielectrics with differing conductive values.

**Figure 3.1:**
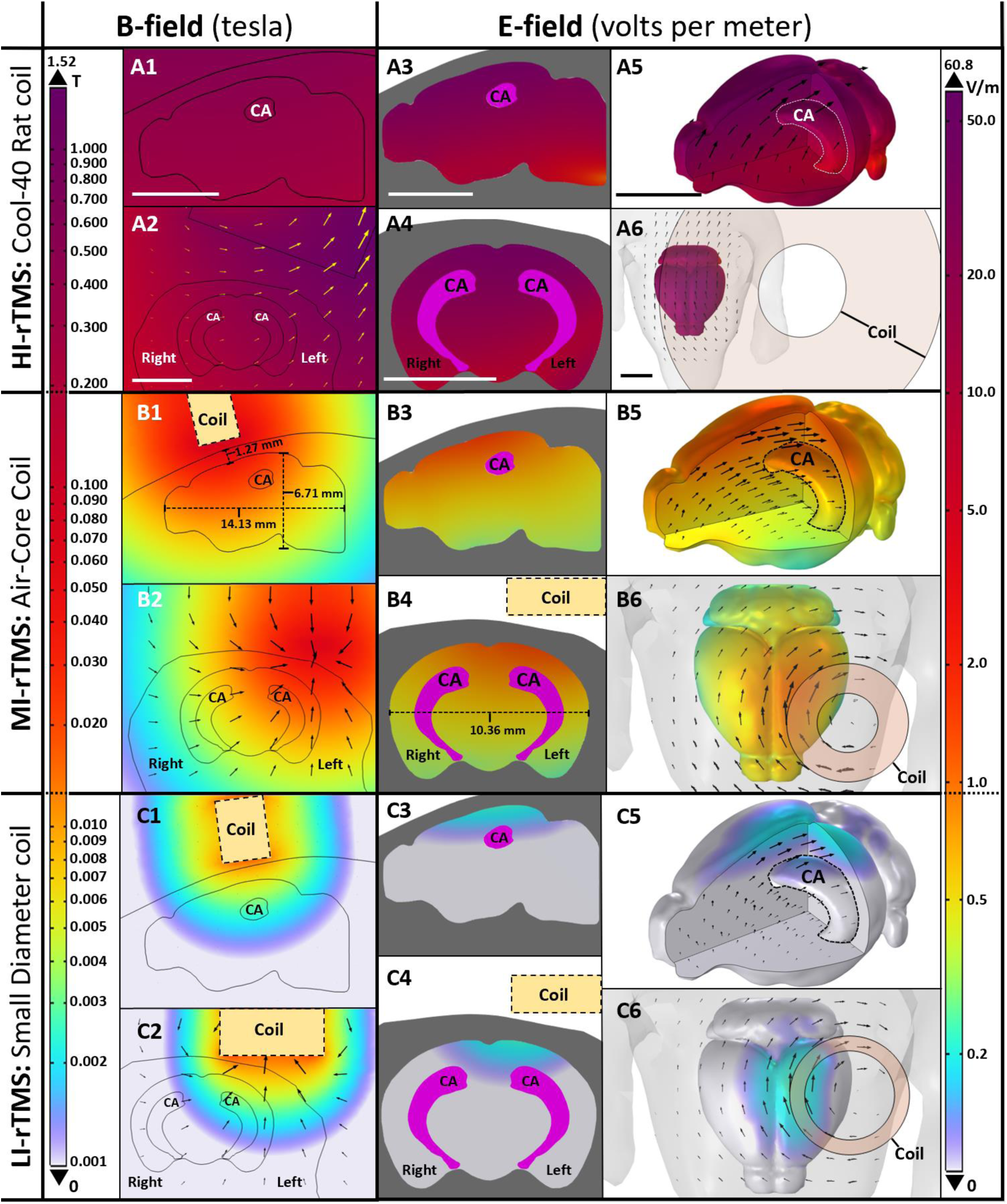
The 12 mT LI-TMS Small Diameter, 90 mT MI-TMS Air-Core, and 1.20 T HI-TMS Cool-40 rat (set to 20% MSO) coils are shown stimulating the mouse brain model over the left hemisphere with logarithmic heat maps shown for both the B-field and E-field. With A1, B1 and C1 and A2, B2 and C2 showing sagittal and coronal planes respectively of the B-field midway through the left hemisphere. A3, B3 and C3 and A4, B4 and C4 show the sagittal and coronal planes respectively of the E-field midway through the left hemisphere. A5, B5, and C5 show a cut away section of the 3D brain model with the E-field and the hippocampus (CA) visible and black arrows denoting E-field direction. A6, B6 and C6 show the top-down view of the 3D brain with the E-field and coils visible and black arrows denoting E-field direction. Scale bar in first two rows = 10 mm

**Figure 3.2:**
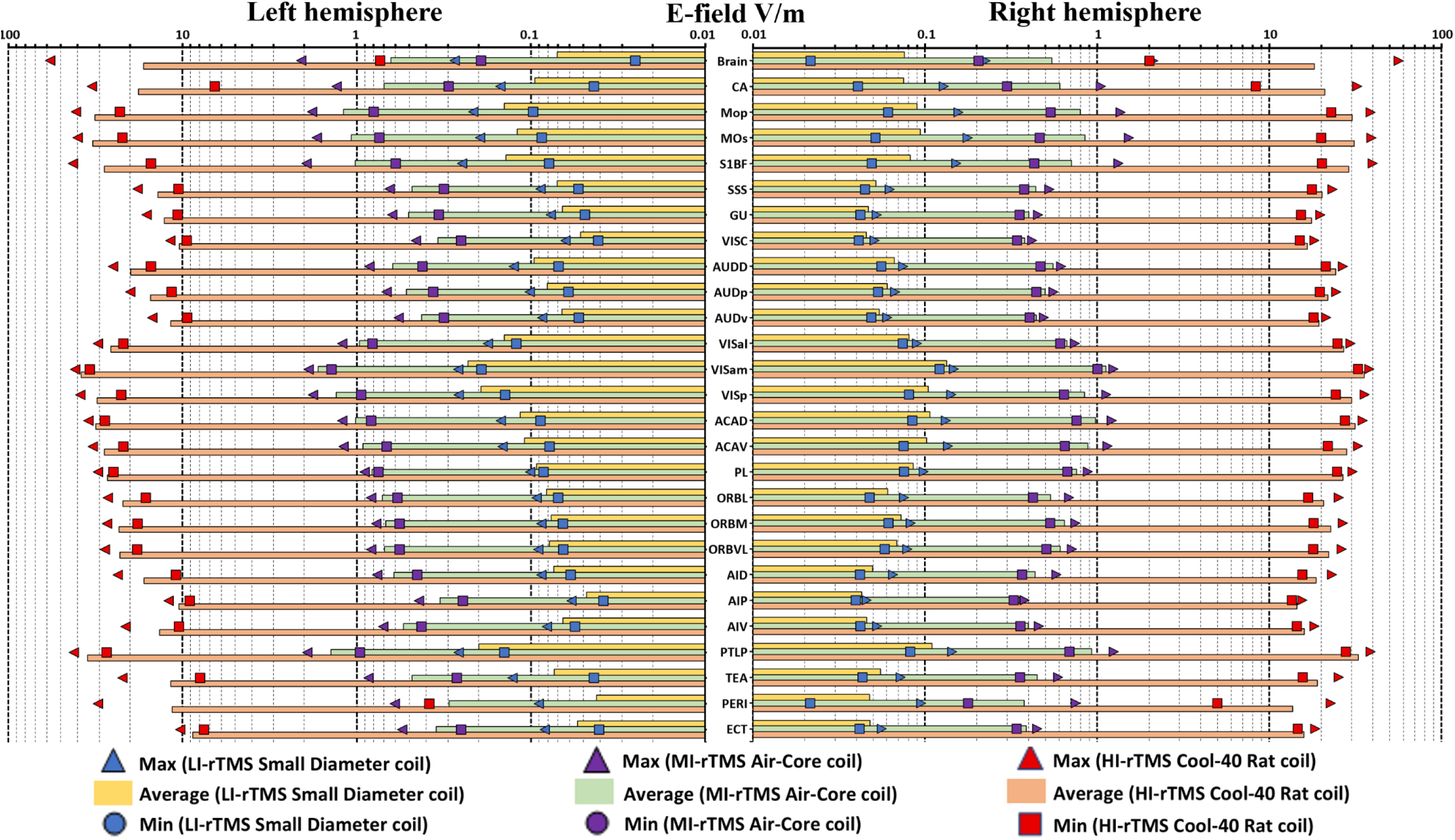
Mouse head regions graphs of E-field values over the left and right brain hemispheres for the 0.012 T LI-TMS Small Diameter, 0.090 T MI-TMS Air-Core, and 1.2 T HI-TMS Cool-40 rodent coils.

**Figure 3.3:**
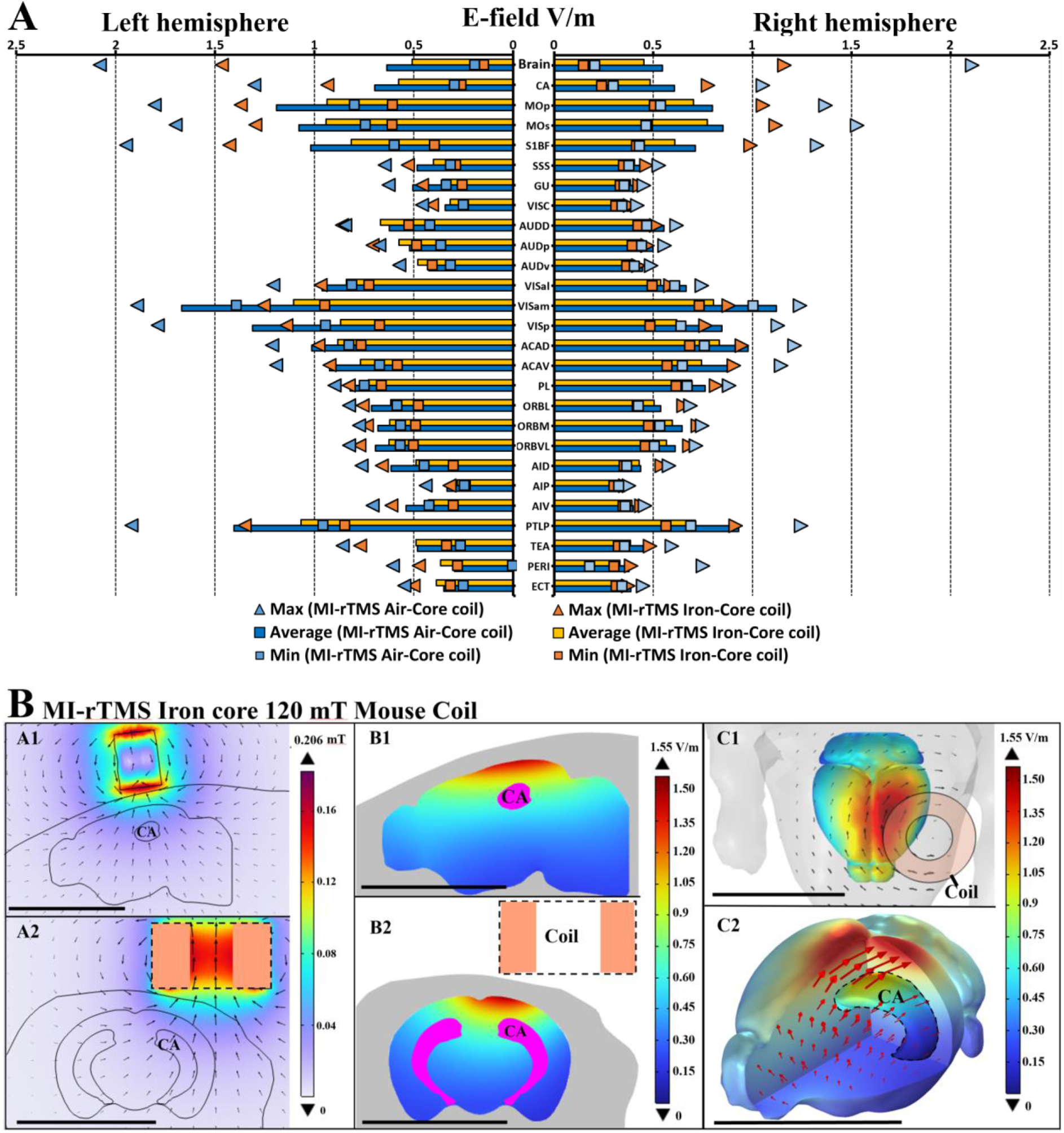
(A) Mouse head regions graphs of E-field values over the left and right brain hemispheres for the 0.120 T MI-TMS Iron-Core rodent coil and the 0.090 T MI-TMS Air-Core. (B) The 0.120 T Iron-Core coil is shown stimulating the mouse brain model over the left hemisphere with peak value heat maps shown for both the B-field and E-field.

### 3.2 B-field and E-field results for rodent coils simulated on the rat head model

The maximum induced B-field and E-field within the rat brain were 6.96×10^-3^ T and 0.144 V/m respectively for the LI-TMS Large Diameter coil, 4.95×10^-2^ T and 0.932 V/m respectively for the MI-TMS Iron-Core coil, and 5.67×10^-1^ T and 97.2 V/m for the HI-TMS D25 Alpha BI coil (See Figure 3.4). Similar to results from the mouse model, a large peak in E-field values from all mouse coils was found at lambda where the most caudal location along midline of the cortex meets the most rostral part of the midbrain forming an angled concave surface (Figure 3.5).

**Figure 3.4:**
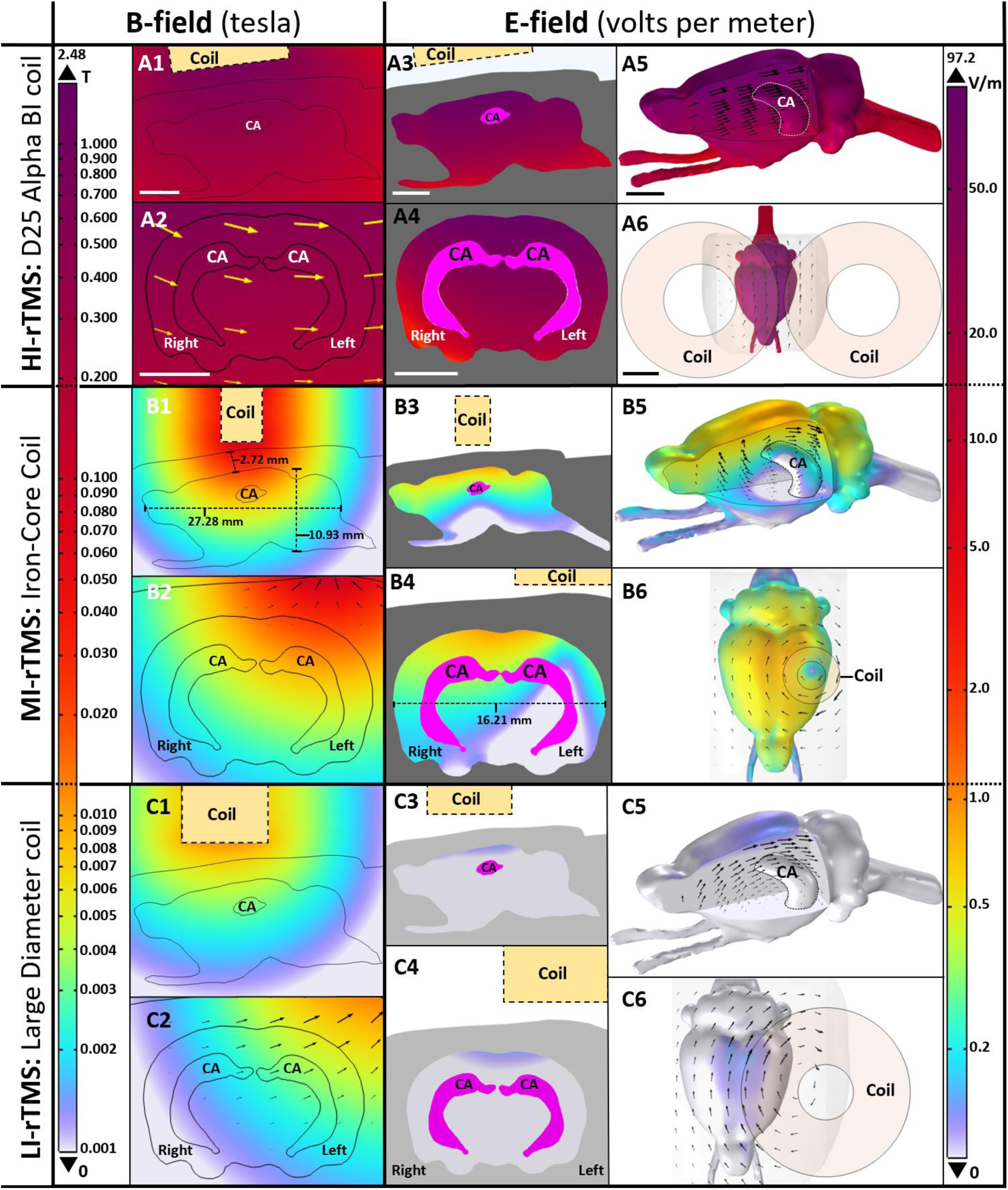
The 13 mT LI-TMS Large Diameter, 120 mT MI-TMS Iron-Core, and 1.16 T HI-TMS D25 Alpha BI (set to 33% MSO) coils are shown stimulating a rat brain model over the left hemisphere with logarithmic heat maps shown for both the B-field and E-field. With A1, B1 and C1 and A2, B2 and C2 showing sagittal and coronal planes respectively of the B-field midway through the left hemisphere. A3-C3 and A4--C4 show the sagittal and coronal planes respectively of the E-field midway through the left hemisphere. A5-C5 show a cut away section of the 3D brain model with the E-field and the hippocampus (CA) visible and black arrows denoting E-field direction. A6-C6 show the top-down view of the 3D brain with the E-field and coils visible and black arrows denoting E-field direction. Scale bar in first two rows = 10 mm

**Figure 3.5:**
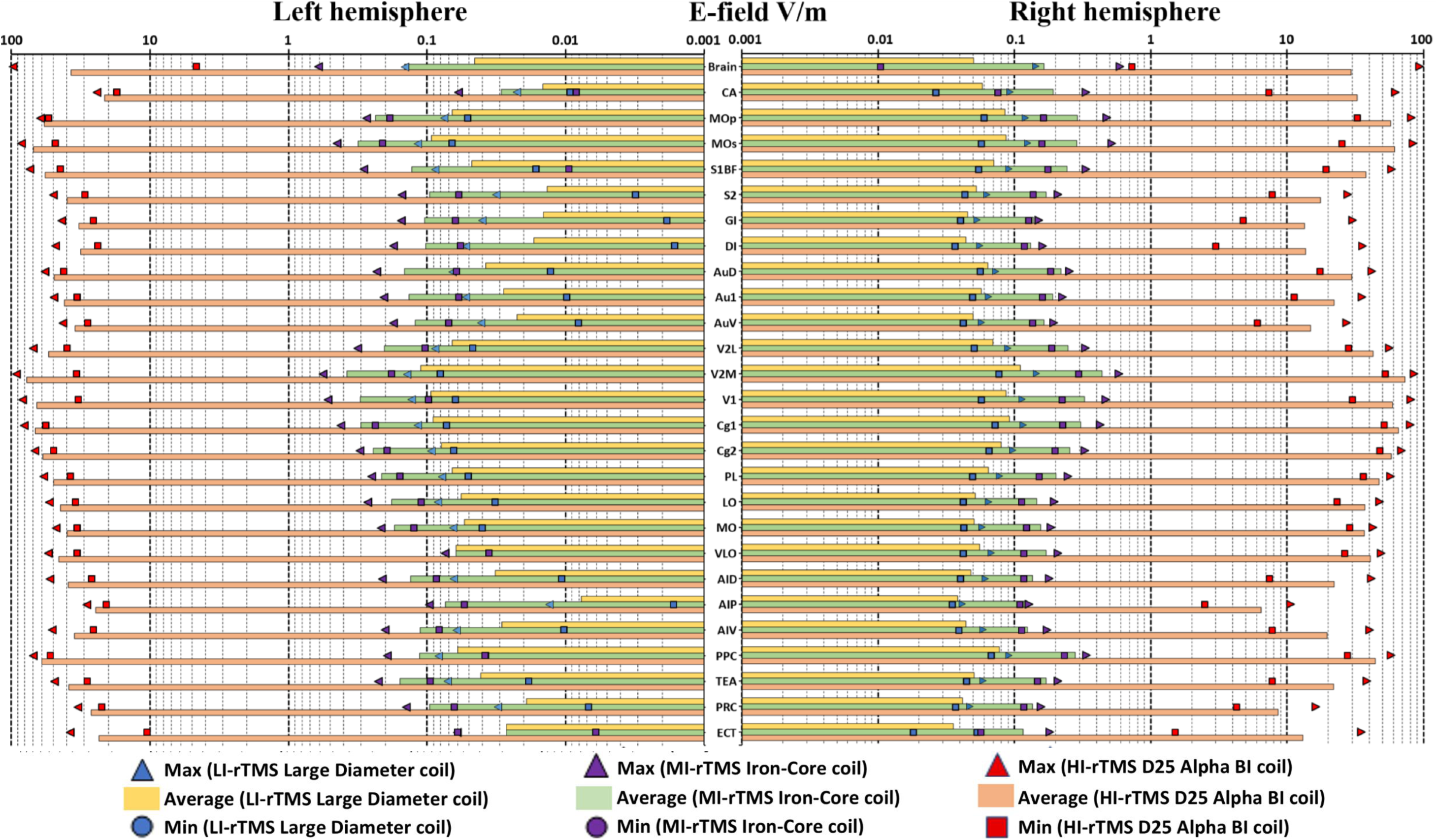
Rat head regions graphs of E-field values over the left and right brain hemispheres for the 0.013 T LI-TMS Large Diameter, 0.090 T MI-TMS Iron-Core, and 1.2 T HI-TMS D25 Alpha BI rodent coils.

### 3.3 B-field and E-field results for clinical TMS coils simulated on a human head model

The maximum induced B-field and E-field within the human brain was 0.282 T and 109.24 V/m respectively for the MC-B70 coil and 0.323 T and 117.74 V/m respectively for the S90 Standard coil, with both coils set to 50% MSO (see Figures 3.6 and 3.7). E-field values for each included brain region can be seen in Figure 3.8. Higher peak values were recorded in sections of geometry with sharp angles, these results were considered modelling artifacts and removed and are discussed in the Supplementary materials section (see Section S.9). Parameters for translational modelling were attained by running parameter sweeps for both coil voltage and distance from the head using the S90 Standard coil and parameters were chosen based on their similarity to rodent E-field modelling results above.

**Figure 3.6:**
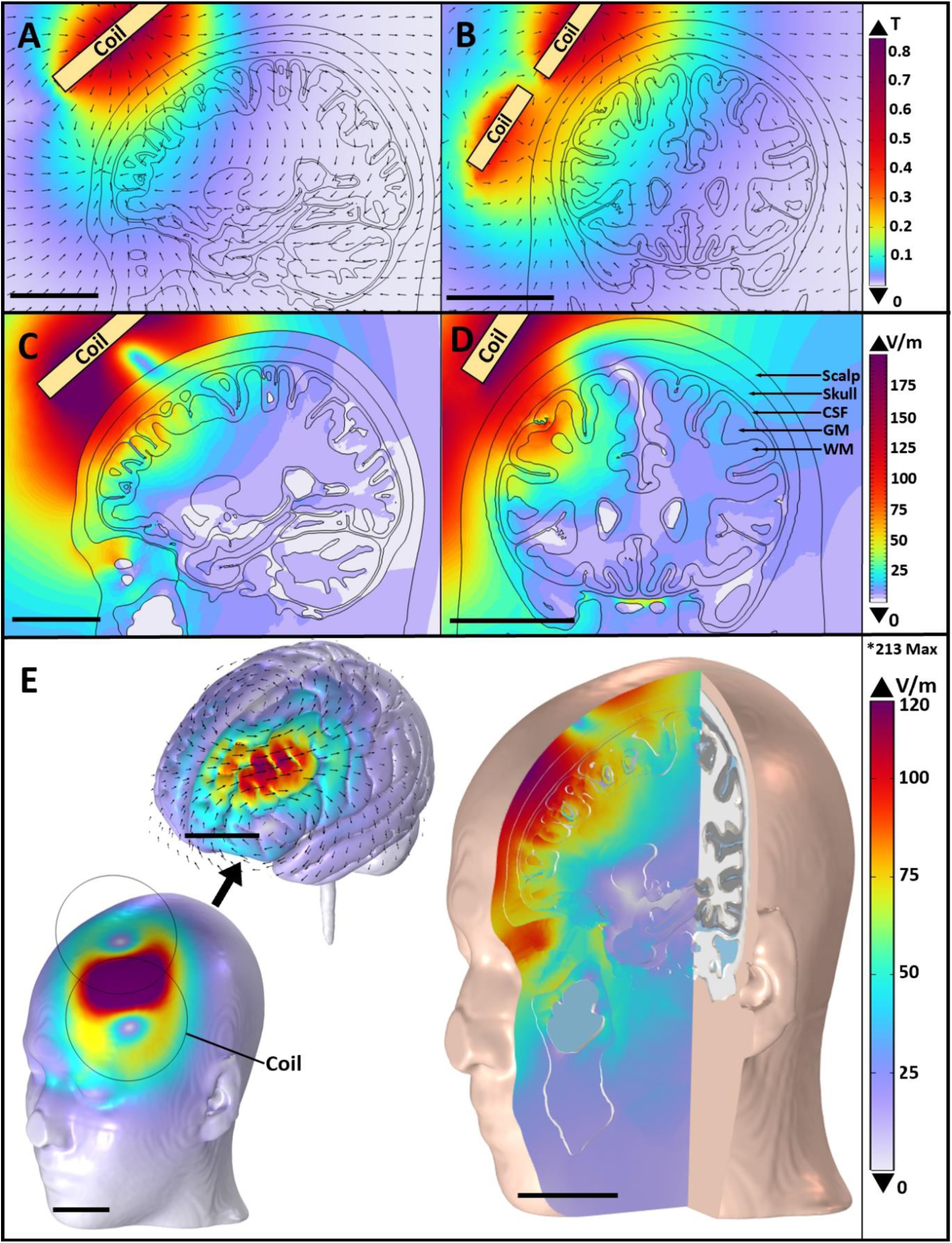
(A) Sagittal and (B) coronal plane views of the human head showing B-field values and (C) Sagittal and (D) coronal plane views, and (E) 3D views of the head, brain and cross section of the head showing E-field values when using the MagVenture MC-B70 coil set to 50% MSO on the MNI152 human head model. Scale bars = 50 mm. *High peak value presented above colour legend (E) was observed in a small location within a sulcus where adjacent sharp GM surface angles are present, as described in Section 3.3.

**Figure 3.7:**
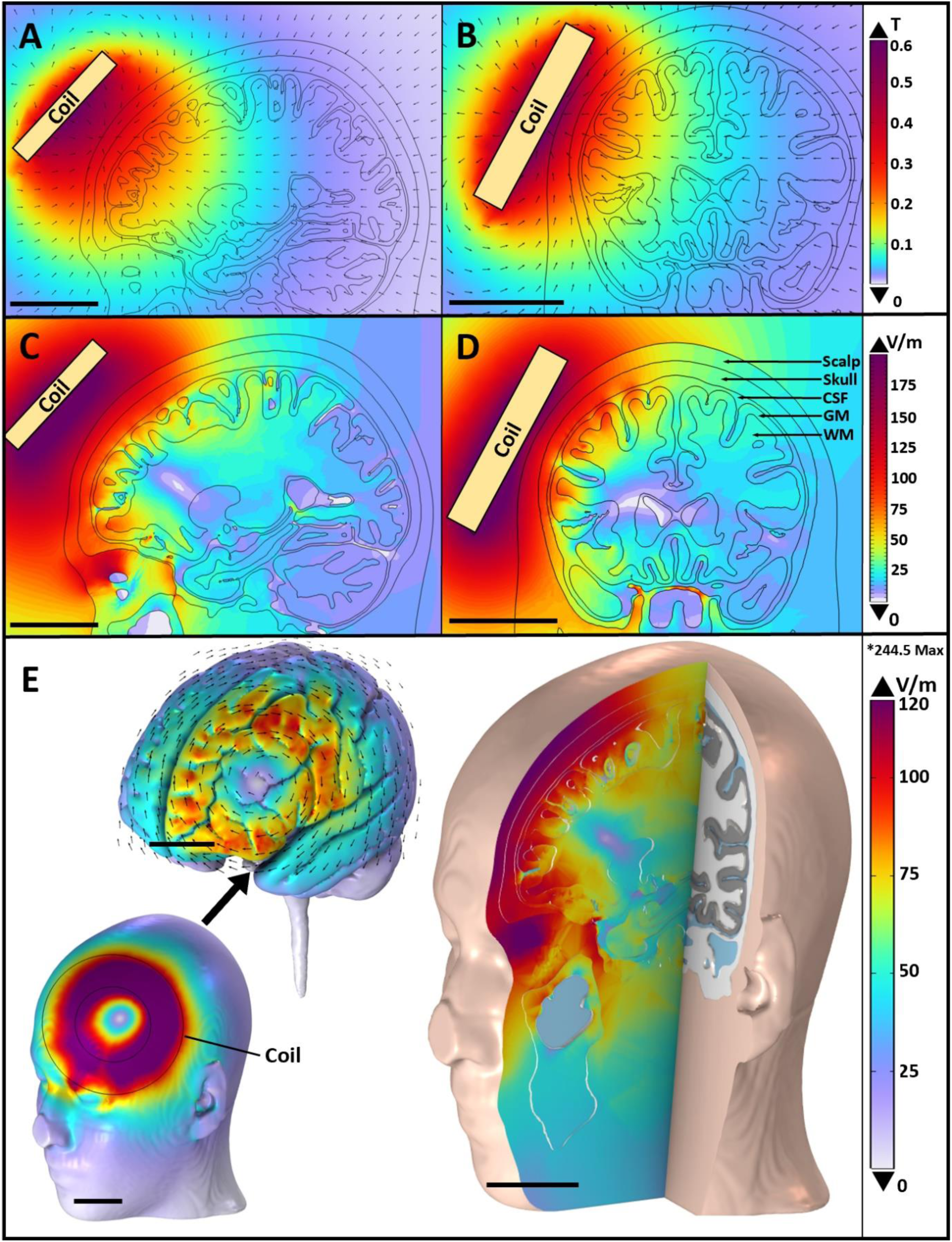
(A) Sagittal and (B) coronal plane views of the human head showing B-field values and (C) Sagittal and (D) coronal plane views, and (E) 3D views of the head, brain and cross section of the head showing E-field values when using the Magstim S90 Standard coil set to 50% MSO on the MNI152 human head model. Scale bars = 50 mm. *High peak value presented above colour legend (E) was observed in a small location within a sulcus where adjacent sharp GM surface angles are present, as described in Section 3.3.

**Figure 3.8:**
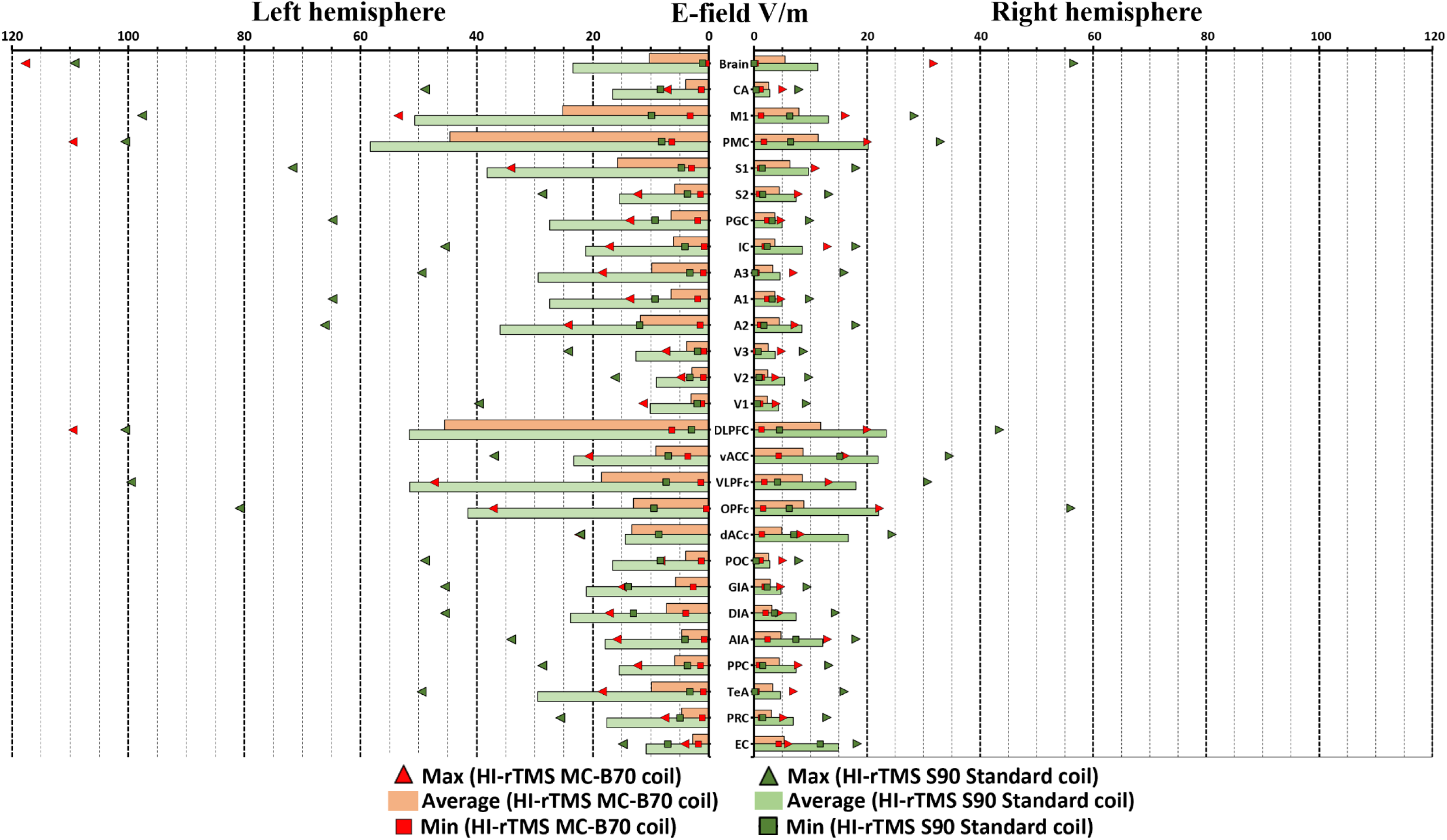
Human head regions graphs of E-field values over the left and right brain hemispheres for the S90 Standard and the MC-B70 coils both set to 50% MSO.

### 3.4 B-field and E-field translational model results for the S90 circular human coil used to replicate rodent TMS studies

The S90 Standard coil was tested on the human head model at varying MSO % settings and distances from the scalp to optimise the replication of rodent LI-TMS and MI-TMS E-field distributions. 2% MSO is a more reliable setting for clinical TMS stimulators due to the propensity for stimulators to shut off intermittently at 1% MSO.

To replicate MI-TMS E-field distributions the S90 coil was set to 1% MSO and 2% MSO with 44 mm extra distance. For replication of LI-TMS E-field distributions the S90 coil was set to 0.1% MSO, 1.0% MSO with 133 mm extra distance, and 2.0% with 177 mm extra distance (See Figures 3.9-3.11). Table 3.1 and Figure 3.12 present the averaged difference in E-field values from all brain regions between the S90 coil translational values and MI-TMS and LI-TMS rodent values.

**Figure 3.9:**
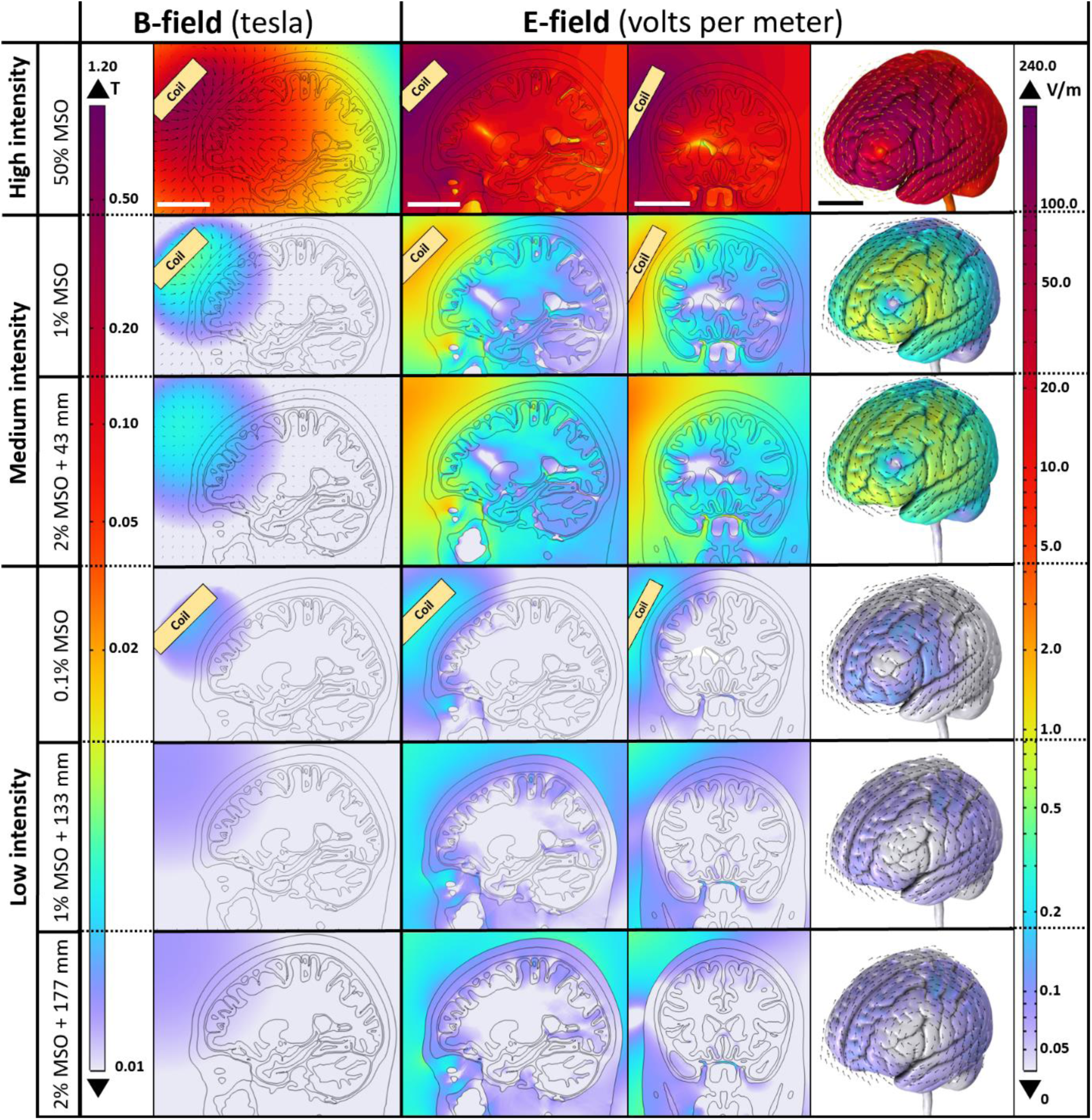
B-field (sagittal plane) and E-field values (sagittal and coronal planes and 3D GM surface) are shown as logarithmic heat maps for the S90 Standard coil stimulating the MNI152 human head model. Moving from the top row to the bottom displayed is the S90 Standard coil at 50% MSO, 1% MSO, 2% MSO with coil placement increased by 43 mm, 0.1% MSO, 1% MSO with coil placement increased by 133 mm, and 2% MSO with coil placement increased by 177 mm. Scale bars along top row = 50 mm

**Figure 3.10:**
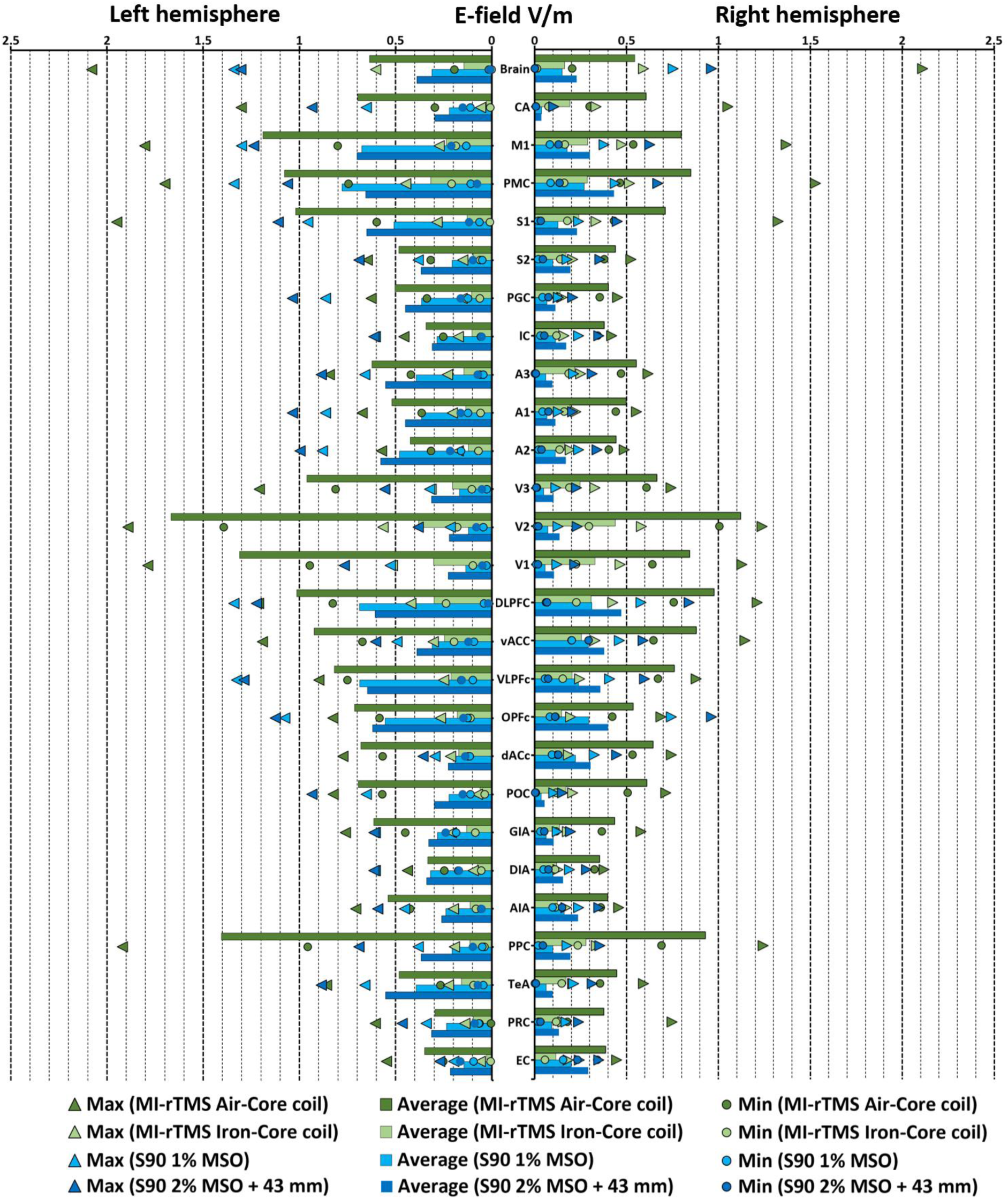
Human head regions graphs of E-field values over the left and right brain hemispheres for the MI-TMS Air-Core, MI-TMS Iron-Core, and S90 Standard coils, with the S90 Standard coil set to 1.0% MSO, and 2% MSO at 44 mm additional distance to replicate E-fields from medium intensity rodent studies.

**Figure 3.11:**
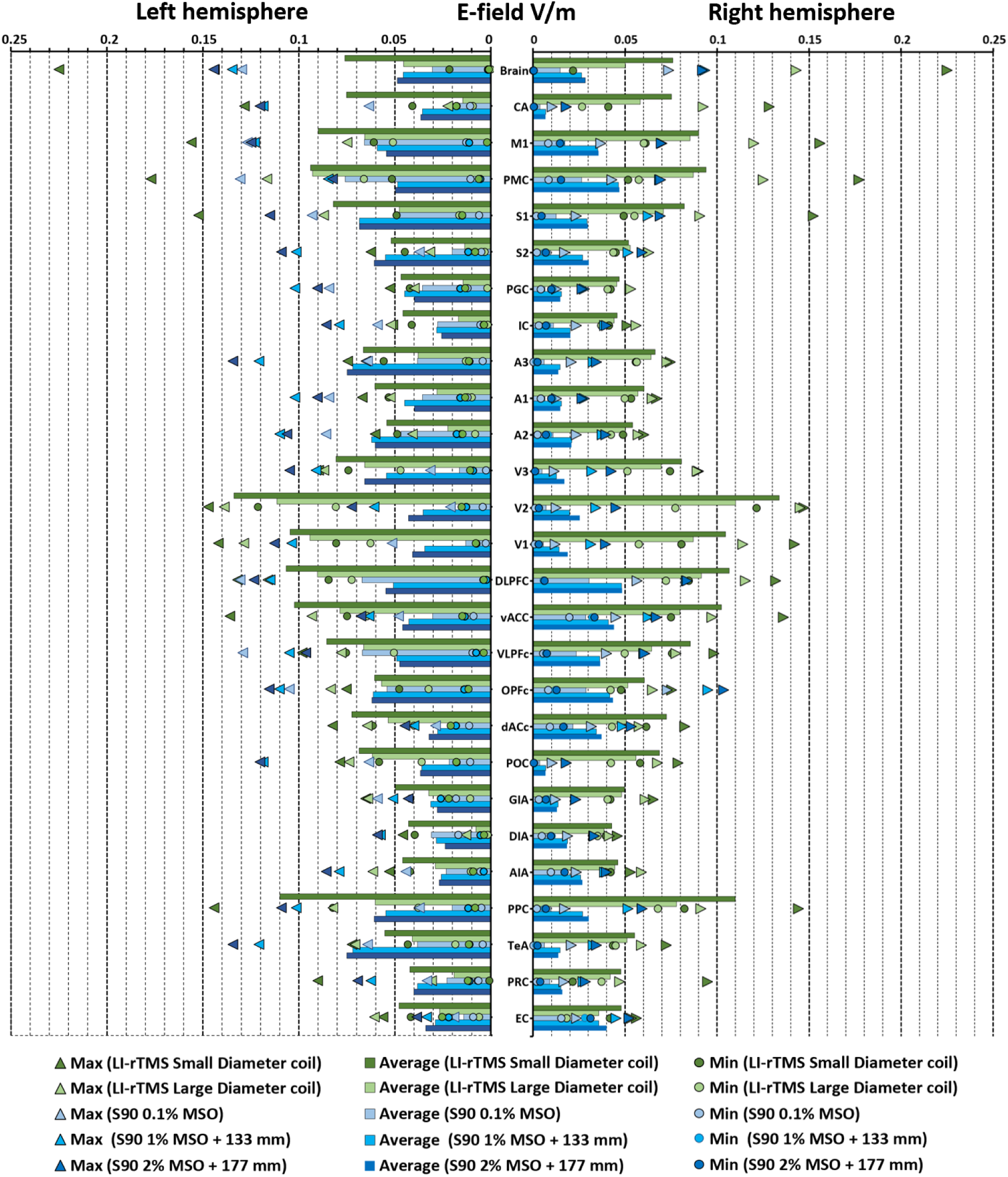
Human head regions graphs of E-field values over the left and right brain hemispheres for the LI-TMS Small Diameter, LI-TMS Large Diameter, and S90 Standard coils, with the S90 Standard coil set to 0.1% MSO, 1% MSO at 133 mm additional distance and 2% MSO at 177 mm additional distance to replicate E-fields from medium intensity rodent studies.

**Figure 3.12:**
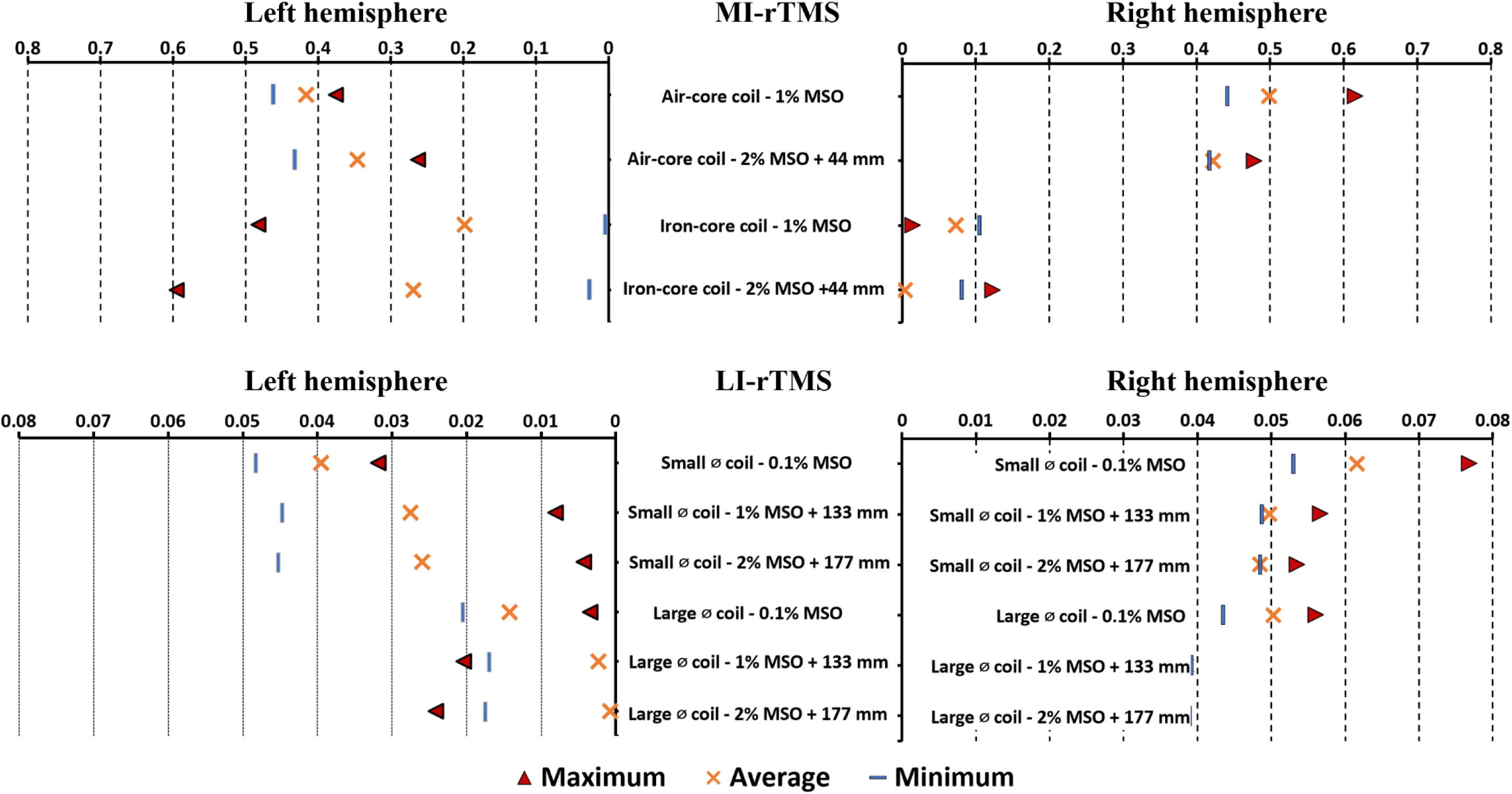
Absolute differences (distance from 0) between rodent model and related translational model counterparts using averaged E-field values across 26 brain regions (maximum, average and minimum) for each combination of LI-TMS and MI-TMS rodent models and their appropriate translational human coil models.

**Table 3.1:**
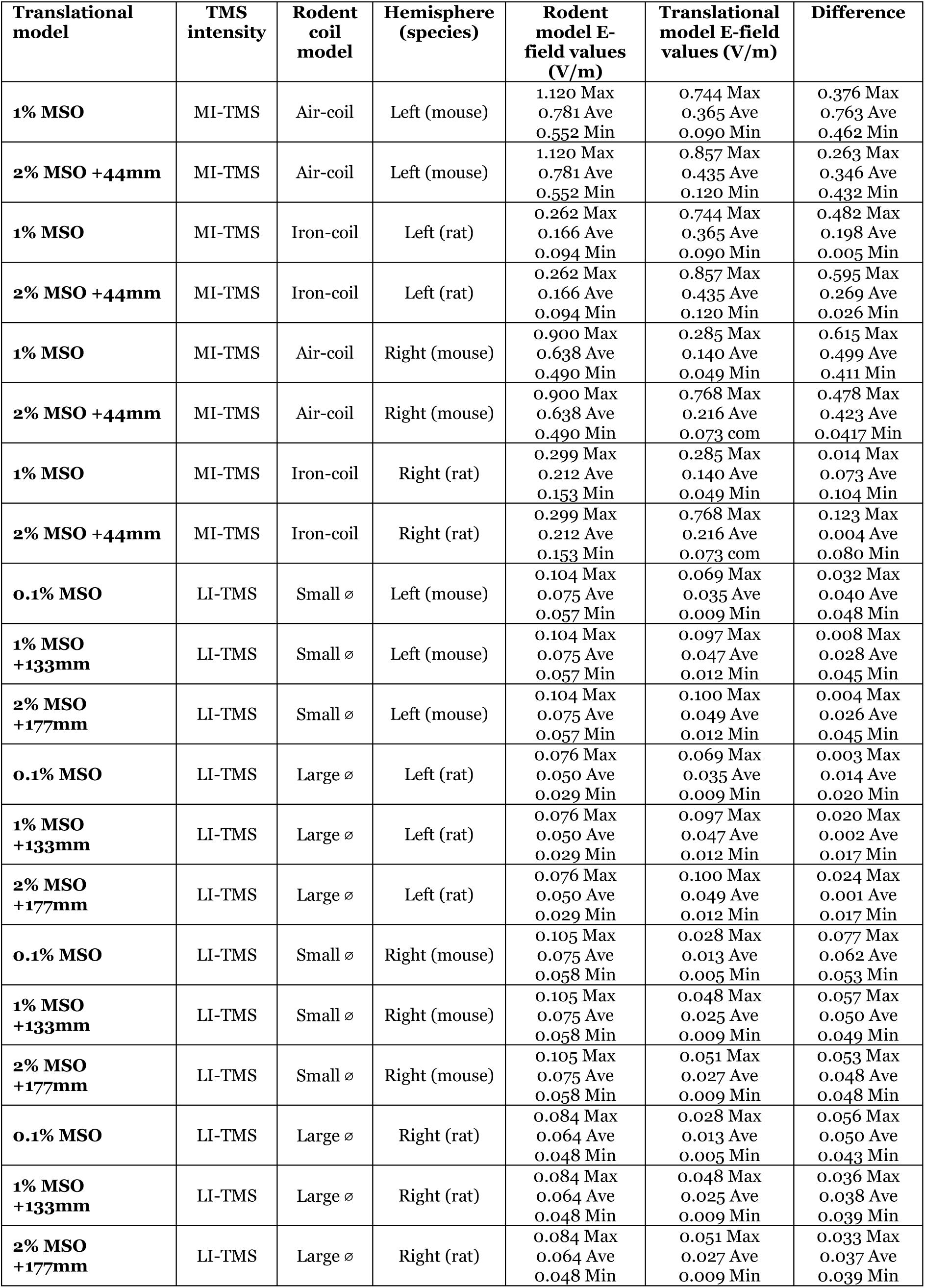
Averaged E-field values across 26 brain regions (maximum, average and minimum) for LI-TMS and MI-TMS rodent and translational human coil models. The difference between rodent models and their appropriate translational model counterparts is presented in the right-most column.

| Translational model | TMS intensity | Rodent coil model | Hemisphere (species) | Rodent model E-field values (V/m) | Translational model E-field values (V/m) | Difference |
| --- | --- | --- | --- | --- | --- | --- |
| **1% MSO** | MI-TMS | Air-coil | Left (mouse) | 1.120 Max<br>0.781 Ave<br>0.552 Min | 0.744 Max<br>0.365 Ave<br>0.090 Min | 0.376 Max<br>0.763 Ave<br>0.462 Min |
| **2% MSO +44mm** | MI-TMS | Air-coil | Left (mouse) | 1.120 Max<br>0.781 Ave<br>0.552 Min | 0.857 Max<br>0.435 Ave<br>0.120 Min | 0.263 Max<br>0.346 Ave<br>0.432 Min |
| **1% MSO** | MI-TMS | Iron-coil | Left (rat) | 0.262 Max<br>0.166 Ave<br>0.094 Min | 0.744 Max<br>0.365 Ave<br>0.090 Min | 0.482 Max<br>0.198 Ave<br>0.005 Min |
| **2% MSO +44mm** | MI-TMS | Iron-coil | Left (rat) | 0.262 Max<br>0.166 Ave<br>0.094 Min | 0.857 Max<br>0.435 Ave<br>0.120 Min | 0.595 Max<br>0.269 Ave<br>0.026 Min |
| **1% MSO** | MI-TMS | Air-coil | Right (mouse) | 0.900 Max<br>0.638 Ave<br>0.490 Min | 0.285 Max<br>0.140 Ave<br>0.049 Min | 0.615 Max<br>0.499 Ave<br>0.411 Min |
| **2% MSO +44mm** | MI-TMS | Air-coil | Right (mouse) | 0.900 Max<br>0.638 Ave<br>0.490 Min | 0.768 Max<br>0.216 Ave<br>0.073 com | 0.478 Max<br>0.423 Ave<br>0.0417 Min |
| **1% MSO** | MI-TMS | Iron-coil | Right (rat) | 0.299 Max<br>0.212 Ave<br>0.153 Min | 0.285 Max<br>0.140 Ave<br>0.049 Min | 0.014 Max<br>0.073 Ave<br>0.104 Min |
| **2% MSO +44mm** | MI-TMS | Iron-coil | Right (rat) | 0.299 Max<br>0.212 Ave<br>0.153 Min | 0.768 Max<br>0.216 Ave<br>0.073 com | 0.123 Max<br>0.004 Ave<br>0.080 Min |
| **0.1% MSO** | LI-TMS | Small ∅ | Left (mouse) | 0.104 Max<br>0.075 Ave<br>0.057 Min | 0.069 Max<br>0.035 Ave<br>0.009 Min | 0.032 Max<br>0.040 Ave<br>0.048 Min |
| **1% MSO +133mm** | LI-TMS | Small ∅ | Left (mouse) | 0.104 Max<br>0.075 Ave<br>0.057 Min | 0.097 Max<br>0.047 Ave<br>0.012 Min | 0.008 Max<br>0.028 Ave<br>0.045 Min |
| **2% MSO +177mm** | LI-TMS | Small ∅ | Left (mouse) | 0.104 Max<br>0.075 Ave<br>0.057 Min | 0.100 Max<br>0.049 Ave<br>0.012 Min | 0.004 Max<br>0.026 Ave<br>0.045 Min |
| **0.1% MSO** | LI-TMS | Large ∅ | Left (rat) | 0.076 Max<br>0.050 Ave<br>0.029 Min | 0.069 Max<br>0.035 Ave<br>0.009 Min | 0.003 Max<br>0.014 Ave<br>0.020 Min |
| **1% MSO +133mm** | LI-TMS | Large ∅ | Left (rat) | 0.076 Max<br>0.050 Ave<br>0.029 Min | 0.097 Max<br>0.047 Ave<br>0.012 Min | 0.020 Max<br>0.002 Ave<br>0.017 Min |
| **2% MSO +177mm** | LI-TMS | Large ∅ | Left (rat) | 0.076 Max<br>0.050 Ave<br>0.029 Min | 0.100 Max<br>0.049 Ave<br>0.012 Min | 0.024 Max<br>0.001 Ave<br>0.017 Min |
| **0.1% MSO** | LI-TMS | Small ∅ | Right (mouse) | 0.105 Max<br>0.075 Ave<br>0.058 Min | 0.028 Max<br>0.013 Ave<br>0.005 Min | 0.077 Max<br>0.062 Ave<br>0.053 Min |
| **1% MSO +133mm** | LI-TMS | Small ∅ | Right (mouse) | 0.105 Max<br>0.075 Ave<br>0.058 Min | 0.048 Max<br>0.025 Ave<br>0.009 Min | 0.057 Max<br>0.050 Ave<br>0.049 Min |
| **2% MSO +177mm** | LI-TMS | Small ∅ | Right (mouse) | 0.105 Max<br>0.075 Ave<br>0.058 Min | 0.051 Max<br>0.027 Ave<br>0.009 Min | 0.053 Max<br>0.048 Ave<br>0.048 Min |
| **0.1% MSO** | LI-TMS | Large ∅ | Right (rat) | 0.084 Max<br>0.064 Ave<br>0.048 Min | 0.028 Max<br>0.013 Ave<br>0.005 Min | 0.056 Max<br>0.050 Ave<br>0.043 Min |
| **1% MSO +133mm** | LI-TMS | Large ∅ | Right (rat) | 0.084 Max<br>0.064 Ave<br>0.048 Min | 0.048 Max<br>0.025 Ave<br>0.009 Min | 0.036 Max<br>0.038 Ave<br>0.039 Min |
| **2% MSO +177mm** | LI-TMS | Large ∅ | Right (rat) | 0.084 Max<br>0.064 Ave<br>0.048 Min | 0.051 Max<br>0.027 Ave<br>0.009 Min | 0.033 Max<br>0.037 Ave<br>0.039 Min |

### 3.5 Comparison of E-field distributions across the cortex between human and rat model

To compare the differences in E-field distributions between the heavily folded human cortex and the relatively smooth rodent cortex the E-field distribution across an arc length of 10 mm of cortex was measured parallel to the base of the coil (Figure 3.13). The simulated coil used for both models was the MC-B70 at 50% held at the same distance from both cortex surfaces.

**Figure 3.13:**
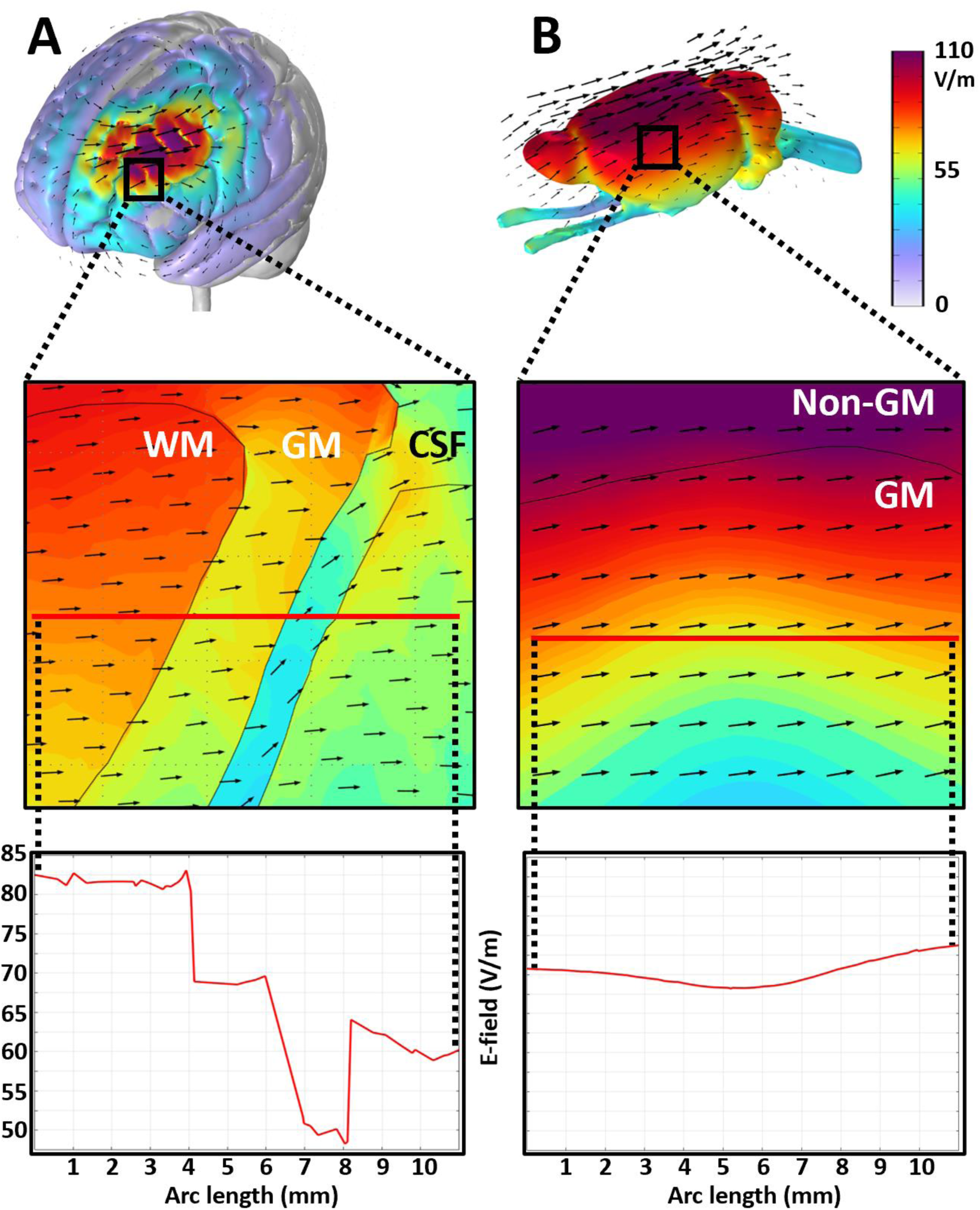
E-field magnitude (V/m) across a 10 mm line parallel with the base of the MC- B70 Magventure coil for both the (A) human and (B) rat brains at a similar depth. Scale bar = 10 mm.

## 4. Discussion

In this study, region-based E-field distributions were simulated using different intensities of TMS applied in previous studies in mouse and rat models. We then provided a pipeline for translation of these TMS parameters into humans. Simulated LI-TMS and MI-TMS in rodents described E-field values approximately three and two orders of magnitude below typical clinical TMS values respectively. Using a commercially available 90 mm circular clinical TMS coil we reproduced LI-TMS and MI-TMS rodent E-field values in a human head by a combination of reducing the MSO% and increasing coil distance from the scalp. As expected, coil positioning was a significant factor in optimising the translation of E-fields, however, results were also affected by species-based differences in brain region morphology, brain surface orientation, coil geometry, and dielectric properties of tissue layers.

### 4.1 Differences in E-fields between mouse and rat TMS

Both LI-TMS and MI-TMS rodent coils induced higher E-field values and larger relative distributions in the mouse than the rat brain models despite similar stimulation parameters. Differences in brain size and the coil-cortical surface distance between species likely result in deeper E-field penetration in mouse compared to rat brain. In the mouse model, the MI-TMS iron-core coil produced similar B-field results to the MI-TMS air-core coil but unexpectedly the E-field distribution was lower in the presence of the iron-core. Section S.3 (see Supplementary materials) resolved this finding as due to the nonlinear behaviour of the iron core coil at the given parameters.

Head and brain size are key factors impacting TMS E-field strength with modelling by Alekseichuk, Mantell et al. (2019) previously detailing a two-term exponential increase followed by a decrease in E-field intensity as head size increased while keeping coil size static. Our paper’s translational model details experimentally and clinically relevant coil sizes specific to each species and as such can be easily used to mitigate differences in E-field distribution due to differences in head size. Furthermore, cropped TMS models are subject to a decrease in E-field intensity (Mantell, Perera et al. 2023), as such whole body rodent and whole head human FEM models are necessary to simulate a realistic E-field distribution, such as those produced in this study (see Supplementary Sections S.8 and S.9).

Our previous FEM model of rodent TMS, used by Tang, Bennett et al. (2021) estimated E-field cortex values to be one magnitude larger than this study’s model. The discrepancy in E-field magnitude is likely due to updated geometries, physics interface parameters and coil stimulation parameters. Supplementary Table S7 contains notes on modelling aspects we consider most integral for accuracy of results. These factors are specific to COMSOL models. Supplementary Section S.9.3 provides a visual representation of how the above issues impact E-field distribution.

### 4.2 Coil positioning impacts E-field distribution differently in rodent and human brains

E-field values were recorded across 26 brain regions for a mouse, rat, and human model, using eight different coils. Differences in size and shape of brain regions between species made translating maximum and minimum values difficult and ultimately, average values were easier to keep consistent across species in multiple brain regions. E-field values varied more between species in deeper structures due to differences in brain volume (Weissman, Epstein et al. 1992) and the non-linear decrease in E-field magnitude with distance from the coil base, as described by the Biot-Savart law:

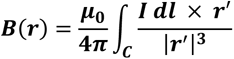

Conversely, replicating E-field values in deeper brain regions between species produces a discrepancy in superficial regions (Deng, Lisanby et al. 2014). A more specific break down of how TMS induced E-fields are governed by Faraday’s law and the Biot-Savart Law is given in Supplementary Section

Due to the small size of the rodent head, our results suggest that HI-TMS directly stimulates deep brain regions as well as the superficial ones. This is readily apparent when observing the greater difference in translated hippocampal E-field values between rodent and human results when compared to any of the more superficial brain region values. There is also a larger disparity of translated E-field values between species in the right hemisphere due to the relatively larger head of humans. This may compromise translation of results from pre-clinical studies because direct TMS stimulation of subcortical regions does not occur in humans. Moreover, direct stimulation of deep brain regions may not be necessary: targeting cortical areas leads to the modulation of functional connectivity with deeper brain structures and may underpin therapeutic outcomes in treatment-resistant depression (George, Nahas et al. 2000); (Klomjai, Katz et al. 2015, Kang, Lee et al. 2016). The relatively shallow penetration of lower intensity TMS in the rodent brain may therefore be more translationally relevant compared to HI-TMS, particularly since connectomes across brains of mammalian species are conserved relative to taxonomic distance (Jonckers, Van Audekerke et al. 2011, Bergmann, Zur et al. 2016, Suárez, Paolino et al. 2018, Goulas, Majka et al. 2019, Assaf, Bouznach et al. 2020, Suarez, Yovel et al. 2022).

### 4.3 Difference in E-field distribution between rodents and humans due to brain geometry, cortical folding, and coil geometry

TMS induced E-fields run parallel to the coil current within the cortex, which is heavily folded in humans and relatively smooth in rodents. This difference, along with variations in head size between species, makes it challenging to replicate similar E-fields in corresponding brain regions. Brain folds vary the depth of brain regions along the cortical surface making translating E-field values across multiple brain regions difficult, for instance, the insula cortex in humans lies deep within a fold but on the surface in rodent brains.

Consistent with findings from Thielscher, Opitz et al. (2011), the complex gyral folding patterns found in FEM human brain models caused local E-field disruptions where the E-field direction crossed between CSF, GM and WM boundaries. E-fields oriented perpendicular to gyral surfaces produced increased E-field intensity in the GM. Furthermore, as observed by Miranda, Hallett et al. (2003), a capacitive effect was present at tissue boundaries resulting in an increase of charge relative to the difference in conductivity within the tissue of lower conductivity (see Figure 3.12). Both these effects (seen in Figure 3.12) are almost absent in rodent brains but significantly contribute E-field distributions in the human brain.

Coil geometry can be used to alter the E-field distribution, which can help translate TMS studies between species. Figure of 8 TMS coils are useful for increasing E-field focus and targeting specific regions. However, for parameters that have first been identified in rodent studies (Cullen, Senesi et al. 2019), circular coils may be better able to replicate the preclinically validated focality and intensity conditions as has been implemented in a recent clinical trial of MI-rTMS in Multiple Sclerosis (Makowiecki, Stevens et al. 2022). Regardless, for any TMS coil simple steps such as changing %MSO and placement distance can refine focality, depth and intensity parameters to match those applied in rodents (see Figure 3.8). For example, our human model using 2% MSO combined with distance reproduced rodent E-field distributions more accurately than the <1% MSO applied directly above the head.

### 4.4: Study limitations

Dielectric values from human tissues are well documented (Gabriel, Lau et al. 1996, Hasgall, Di Gennaro et al. 2018) and, as in this study, are routinely applied to rodent FEM TMS models. However, cell morphology and density vary between mammalian species proportionally to differences in brain size (Herculano-Houzel, Manger et al. 2014). While bovine, ovine, and porcine dielectric values are similar to human dielectric values (Matković, Kordić et al. 2022), rodent values are likely to differ. Detailed recordings of rodent tissue dielectric values at various stimulation frequencies are not currently available and would improve accuracy and better guide translational modelling efforts.

The impact of conductivity, permeability, and permittivity (dielectric values) on the E-field distribution was investigated and conductivity was observed to be the determinant factor for E-field distribution with permittivity only impacting results at artificially larger value differences of approximately 10^4, between tissues. However, as previously acknowledged (Greiner 2012, Cvetković, Poljak et al. 2015, Gaugain, Quéguiner et al. 2023, Wang, Peterchev et al. 2024), further increases in geometry complexity and the future implementation of multiscale modelling led to permittivity and permeability being potentially impactful and their inclusion is likely required for futureproofing. However, it is also important that modelling artifacts caused by these inclusions in complexity are actively sought and mitigated, although none have been noted in the current study.

Additional modelling artifacts and issues, including the impact of sharp edges in geometry and missing dielectric data, were recognised within this study and avoided where possible, they are described in detail in Supplementary Section 9 along with methods for mitigating their impact . FEM modelling artifacts for TMS described in this work primarily emerged due to the necessary simplification of real-world geometry along with how COMSOL’s equations describe the interaction of electromagnetic fields with this geometry.

### 4.5: Future directions

This is the first study to provide a translational model for replicating TMS-derived E-fields validated in rodent models in the human brain. However it is clear that our model does not capture the nuances of human or rodent neurobiological responses to TMS: we do not resolve how LI-TMS and MI-TMS E-field distributions interact directly with biomolecules, neurons, glia or cell-scale geometries and we do not account for differences in functional neuroanatomy between species. Rather, we aimed to provide a model that will facilitate delivery of precisely defined electric fields in a range of head sizes across species, which is a key requirement for being able to study the biological mechanisms of TMS and develop robust therapeutic protocols.

A multiscale model, inclusive of a Hodgkin-Huxley model (e.g. NEUON (Hines and Carnevale 1997))has the potential to describe TMS induced changes at the molecular, cell, and neural network scale. This is of particular interest in LI-TMS and MI-TMS research where subthreshold plasticity seems to be guided by intricate cellular and molecular phenomena (Moretti and Rodger 2022). For example, a multiscale modelling toolbox developed by Shirinpour, Hananeia et al. (2021) describes TMS induced calcium accumulation, a phenomena induced experimentally using LI-rTMS (Grehl, Viola et al. 2015, Clarke, Penrose et al. 2017).

Cell level modelling would describe if the E-field normal component derived from lower intensity TMS has an impact on subthreshold plasticity, similar to how modelling by Aberra, Wang et al. (2020) described the E-field normal component of clinical intensity TMS being independent of action potential generation. This information could contribute to a better understanding of the therapeutic potential of harnessing subthreshold plasticity in neurological conditions. Furthermore, neural orientation differs markedly between rodents and humans based on the presence or absence of gyri, which presents an issue of unknown significance in the replication of E-fields across species. Multiscale modelling efforts into investigating the impact of subthreshold E-field normal components would elucidate this point and could help contribute to refining TMS techniques for specific neural effects. Multiscale models can also help elucidate whether particular cell types, cell alignment, geometry, molecular phenomena, and myelination effect the E-field distribution (Siebner, Funke et al. 2022). However, there are challenges and limitations with the implementation of multiscale models and with increasing complexity: significant computational power is required, and the likelihood of error or misinterpretation is high, particularly with the merging of two different forms of modelling and mathematics (FEM and Hodgkin-Huxley modelling).

Avoidance of coil misplacement and variability during future translational work using this paper’s model will require attention as accuracy is highly dependent on spatial replication of experimental setups and the relatively lower E-field penetration of LI-rTMS and MI-rTMS could potentially lead to a higher level of mis-targeting and intra-individual variability than with clinical TMS (Koehler, Kammer et al. 2023).

Specific research questions pertinent to the future accuracy and utility of the translational model include: Firstly, how can the values from an FEM derived TMS induced E-field, and its associated aspects, be applied to a Hodgkin Huxley model accurately and with validation for incorporation of a multiscale model to better investigate cellular and molecular phenomena of TMS and is NEURON’s extracellular function able to accurately describe a TMS E-field? Secondly, what are dielectric values of rodent brain tissues and does brain anisotropy alter the electric field distributions? Thirdly, does variation in rodent head size and skull thickness impact low intensity field distribution as has been demonstrated in humans for high intensity rTMS? Fourthly, our model can be used to identify biological markers in animal models that can indicate neuronal or glial cell responsiveness to different intensities of TMS. This would not only provide a real-world validation of our model, but could potentially translate into humans (eg via MRI or other imaging modality) to further refine TMS applications and understanding.

The translational model offers an ethically advantageous predictive method for E-field distribution, particularly once experimentally validated, by lowering the requirement of explorative studies thereby reducing the number of animal models in LI-rTMS studies and as a result expediting development of novel TMS treatments. However, failure of experimental validation will still provide valuable insight into model limitations, guiding future efforts.

### 4.6: Conclusion

This study produced the first known FEM model for translating LI-TMS and MI-TMS induced E-fields from rodents across multiple brain regions into humans. This was achieved with a current commercial stimulator (S90 Standard coil) in combination with considered coil placement. This translational model will be useful in designing stimulation parameters in humans based on rodent studies, furthermore, it can be used to model clinical TMS parameters in rodents and to help produce custom coil designs.

Typically, B-field values are used to describe TMS coils and sometimes TMS intensity in the brain stimulation field. Our models suggest E-field intensity both in the whole brain and in given brain regions of interest provide a more meaningful description of the effects of TMS. This is because, unlike B-fields, simulated E-fields directly interact with neural circuits and the complexity of the E-field distribution (shape, rise and fall time etc) is not intuitive from any given B-field, particularly when brain geometry across species impacts the E-field to a greater degree than the B-field. Furthermore, the use of percentage value of motor evoked potential threshold (regularly used in TMS research) may not be relevant for circuits and brain regions that are different to the motor cortex (e.g., cerebellum). Bridging the gap between animal model studies of novel treatments for TMS and clinical trials is an important and complex undertaking with work by Makowiecki, Stevens et al. (2022) being a notable recent example. However, with the increase in open-source models, such as the present study and other software (e.g., SimNIBS), providing simulated E-field values in addition to other stimulation parameters translation should become more feasible.

## CrediT authorship contribution statement

**Samuel J. Bolland:** - Conceptualization, data collection, analysis, and interpretation, Writing first draft of manuscript, review, and editing

**Maxim Goryachev:** - Data collection analysis and interpretation, Writing first draft of manuscript, review and editing, Supervision

**Alex Opitz:** Data interpretation, Writing – review and editing

**Alex D. Tang:** - Conceptualization, data interpretation, Writing – review & editing, Supervision, Project administration, Funding acquisition.

**Jennifer Rodger:** Conceptualization, data interpretation, Writing – review & editing, Supervision, Project administration, Funding acquisition.

## Abbreviations

TMS: repetitive transcranial magnetic stimulation
TRD: treatment resistant depression
FEM: finite element method
DLPFC: dorsolateral prefrontal cortex
GM: grey matter
WM: white matter
CSF: cerebrospinal fluid
MSO: maximum stimulator output

## References

1. Aberra, A. S., et al. (2020). “Simulation of transcranial magnetic stimulation in head model with morphologically-realistic cortical neurons.” Brain stimulation 13(1): 175–189.

2. Alekseichuk, I., et al. (2019). “Comparative modeling of transcranial magnetic and electric stimulation in mouse, monkey, and human.” Neuroimage 194: 136–148.

3. Assaf, Y., et al. (2020). “Conservation of brain connectivity and wiring across the mammalian class.” Nature Neuroscience 23(7): 805–808.

4. Bergmann, E., et al. (2016). “The organization of mouse and human cortico-hippocampal networks estimated by intrinsic functional connectivity.” Cerebral Cortex: 1–16.

5. Beynel, L., et al. (2020). “Effects of repetitive transcranial magnetic stimulation on resting-state connectivity: a systematic review.” Neuroimage 211: 116596.

6. Boyer, M., et al. (2022). “In vivo low-intensity magnetic pulses durably alter neocortical neuron excitability and spontaneous activity.” The Journal of Physiology 600(17): 4019–4037.

7. Choung, J. S., et al. (2023). “Development and application of rTMS device to murine model.” Scientific reports 13(1): 5490.

8. Clarke, D., et al. (2017). “Low intensity rTMS has sex-dependent effects on the local response of glia following a penetrating cortical stab injury.” Experimental neurology 295: 233–242.

9. Collins, D. L., et al. (1995). “Automatic 3-D model-based neuroanatomical segmentation.” Human brain mapping 3(3): 190–208.

10. Crowther, L. J., et al. (2014). “Transcranial magnetic stimulation of mouse brain using high-resolution anatomical models.” Journal of Applied Physics 115(17): 17B303.

11. Cullen, C. L., et al. (2019). “Low-intensity transcranial magnetic stimulation promotes the survival and maturation of newborn oligodendrocytes in the adult mouse brain.” Glia 67(8): 1462–1477.

12. Cvetković, M., et al. (2015). “Analysis of transcranial magnetic stimulation based on the surface integral equation formulation.” IEEE Transactions on Biomedical Engineering 62(6): 1535–1545.

13. Davey, K., et al. (2003). “Modeling the effects of electrical conductivity of the head on the induced electric field in the brain during magnetic stimulation.” Clinical Neurophysiology 114(11): 2204–2209.

14. Davey, K. R. and M. Riehl (2006). “Suppressing the surface field during transcranial magnetic stimulation.” IEEE Transactions on Biomedical Engineering 53(2): 190–194.

15. Deng, Z.-D., et al. (2014). “Coil design considerations for deep transcranial magnetic stimulation.” Clinical Neurophysiology 125(6): 1202–1212.

16. Desikan, R. S., et al. (2006). “An automated labeling system for subdividing the human cerebral cortex on MRI scans into gyral based regions of interest.” Neuroimage 31(3): 968–980.

17. Dogdas, B., et al. (2007). “Digimouse: a 3D whole body mouse atlas from CT and cryosection data.” Physics in medicine & biology 52(3): 577.

18. Fitzgerald, P. B., et al. (2006). “A comprehensive review of the effects of rTMS on motor cortical excitability and inhibition.” Clinical Neurophysiology 117(12): 2584–2596.

19. Fitzgerald, P. B., et al. (2009). “A randomized trial of rTMS targeted with MRI based neuro-navigation in treatment-resistant depression.” Neuropsychopharmacology 34(5): 1255–1262.

20. Fonov, V., et al. (2011). “Unbiased average age-appropriate atlases for pediatric studies.” Neuroimage 54(1): 313–327.

21. Fonov, V. S., et al. (2009). “Unbiased nonlinear average age-appropriate brain templates from birth to adulthood.” Neuroimage (47): S102.

22. Foster, K. R. and H. P. Schwan (1986). “Dielectric properties of tissues.” CRC handbook of biological effects of electromagnetic fields: 27–96.

23. Frazier, J. A., et al. (2005). “Structural brain magnetic resonance imaging of limbic and thalamic volumes in pediatric bipolar disorder.” American Journal of Psychiatry 162(7): 1256–1265.

24. Gabriel, C. (1996). Compilation of the dielectric properties of body tissues at RF and microwave frequencies, King’s coll london (United Kingdom) dept of physics.

25. Gabriel, S., et al. (1996). “The dielectric properties of biological tissues: III. Parametric models for the dielectric spectrum of tissues.” Physics in medicine & biology 41(11): 2271.

26. Gaugain, G., et al. (2023). “Quasi-static approximation error of electric field analysis for transcranial current stimulation.” Journal of Neural Engineering 20(1): 016027.

27. George, M. S., et al. (2000). “A controlled trial of daily left prefrontal cortex TMS for treating depression.” Biological psychiatry 48(10): 962–970.

28. Goerzen, D., et al. (2020). “An MRI-derived neuroanatomical atlas of the Fischer 344 rat brain.” Scientific reports 10(1): 6952.

29. Goldstein, J. M., et al. (2007). “Hypothalamic abnormalities in schizophrenia: sex effects and genetic vulnerability.” Biological psychiatry 61(8): 935–945.

30. Goldsworthy, M. R., et al. (2021). “Effects of rTMS on the brain: is there value in variability?” Cortex 139: 43–59.

31. Gomez-Tames, J., et al. (2020). “Review on biophysical modelling and simulation studies for transcranial magnetic stimulation.” Physics in medicine & biology 65(24): 24TR03.

32. Goulas, A., et al. (2019). “A blueprint of mammalian cortical connectomes.” PLoS biology 17(3): e2005346.

33. Grehl, S., et al. (2015). “Cellular and molecular changes to cortical neurons following low intensity repetitive magnetic stimulation at different frequencies.” Brain stimulation 8(1): 114–123.

34. Greiner, W. (2012). Classical electrodynamics, Springer Science & Business Media.

35. Hammelrath, L., et al. (2016). “Morphological maturation of the mouse brain: An in vivo MRI and histology investigation.” Neuroimage 125: 144–152.

36. Hasgall, P., et al. (2015). “IT’IS Database for thermal and electromagnetic parameters of biological tissues.” Version 3.0.

37. Hasgall, P., et al. (2018). “IT’IS Database for thermal and electromagnetic parameters of biological tissues.” Version 4.0.

38. Heath, A., et al. (2018). “Medium-and high-intensity rTMS reduces psychomotor agitation with distinct neurobiologic mechanisms.” Translational psychiatry 8(1): 1–13.

39. Herculano-Houzel, S., et al. (2014). “Brain scaling in mammalian evolution as a consequence of concerted and mosaic changes in numbers of neurons and average neuronal cell size.” Frontiers in neuroanatomy 8: 77.

40. Hines, M. L. and N. T. Carnevale (1997). “The NEURON simulation environment.” Neural computation 9(6): 1179–1209.

41. Jenkinson, M., et al. (2012). “Fsl.” Neuroimage 62(2): 782–790.

42. Jiang, W., et al. (2023). “A C-shaped miniaturized coil for transcranial magnetic stimulation in rodents.” Journal of Neural Engineering 20(2): 026022.

43. Jonckers, E., et al. (2011). “Functional connectivity fMRI of the rodent brain: comparison of functional connectivity networks in rat and mouse.” PloS one 6(4): e18876.

44. Kang, J. I., et al. (2016). “Frontostriatal connectivity changes in major depressive disorder after repetitive transcranial magnetic stimulation: a randomized sham-controlled study.” The Journal of clinical psychiatry 77(9): 7478.

45. Klomjai, W., et al. (2015). “Basic principles of transcranial magnetic stimulation (TMS) and repetitive TMS (rTMS).” Annals of physical and rehabilitation medicine 58(4): 208–213.

46. Koehler, M., et al. (2023). “Quantitative Impact of Coil Misalignment and Misplacement in Transcranial Magnetic Stimulation.” bioRxiv: 2023.2011. 2018.567677.

47. Lein, E. S., et al. (2007). “Genome-wide atlas of gene expression in the adult mouse brain.” Nature 445(7124): 168–176.

48. Madore, M. R., et al. (2021). “Moving back in the brain to drive the field forward: Targeting neurostimulation to different brain regions in animal models of depression and neurodegeneration.” Journal of neuroscience methods 360: 109261.

49. Magstim (2021). “D25 Alpha.” from https://www.magstim.com/us-en/d25-alpha-bi-coil/.

50. Magstim (2021). “S90 Standard Coil.” Retrieved 21/04/2023, from https://www.magstim.com/us-en/s90-standard-coil/.

51. MagVenture (2022). “Cool-40 Rat.” from https://www.magventure.com/us/component/k2/coils/cool-40-rat-coil.

52. MagVenture (2023). “MC-B70.” from https://magventure.com/products/mc-b70/.

53. Makarov, S. N., et al. (2015). “Preliminary upper estimate of peak currents in transcranial magnetic stimulation at distant locations from a TMS coil.” IEEE Transactions on Biomedical Engineering 63(9): 1944–1955.

54. Makowiecki, K., et al. (2018). “Low-intensity repetitive transcranial magnetic stimulation requires concurrent visual system activity to modulate visual evoked potentials in adult mice.” Scientific reports 8(1): 5792.

55. Makowiecki, K., et al. (2014). “Low-intensity repetitive transcranial magnetic stimulation improves abnormal visual cortical circuit topography and upregulates BDNF in mice.” Journal of Neuroscience 34(32): 10780–10792.

56. Makowiecki, K., et al. (2022). “Safety of low-intensity repetitive transcranial magneTic brAin stimUlation foR people living with mUltiple Sclerosis (TAURUS): study protocol for a randomised controlled trial.” Trials 23(1): 1–18.

57. Makris, N., et al. (2006). “Decreased volume of left and total anterior insular lobule in schizophrenia.” Schizophrenia research 83(2-3): 155–171.

58. Manera, A. L., et al. (2020). “CerebrA, registration and manual label correction of Mindboggle-101 atlas for MNI-ICBM152 template.” Scientific Data 7(1): 1–9.

59. Mantell, K. E., et al. (2023). “Anatomical details affect electric field predictions for non-invasive brain stimulation in non-human primates.” Neuroimage 279: 120343.

60. Matković, A., et al. (2022). “Complex Permittivity of Ex-Vivo Human, Bovine and Porcine Brain Tissues in the Microwave Frequency Range.” Diagnostics 12(11): 2580.

61. Mazziotta, J., et al. (2001). “A probabilistic atlas and reference system for the human brain: International Consortium for Brain Mapping (ICBM).” Philosophical Transactions of the Royal Society of London. Series B: Biological Sciences 356(1412): 1293–1322.

62. Miranda, P. C., et al. (2003). “The electric field induced in the brain by magnetic stimulation: a 3-D finite-element analysis of the effect of tissue heterogeneity and anisotropy.” IEEE Transactions on Biomedical Engineering 50(9): 1074–1085.

63. Moretti, J., et al. (2021). “Concurrent LI-rTMS induces changes in c-Fos expression but not behavior during a progressive ratio task with adult ephrin-A2A5-/-mice.” Behavioural brain research 400: 113011.

64. Moretti, J. and J. Rodger (2022). “A little goes a long way: neurobiological effects of low intensity rTMS and implications for mechanisms of rTMS.” Current Research in Neurobiology: 100033.

65. Nowak, K., et al. (2011). “Optimizing a rodent model of Parkinson’s disease for exploring the effects and mechanisms of deep brain stimulation.” Parkinson’s Disease 2011.

66. Papp, E. A., et al. (2014). “Waxholm Space atlas of the Sprague Dawley rat brain.” Neuroimage 97: 374–386.

67. Parthoens, J., et al. (2016). “Performance characterization of an actively cooled repetitive transcranial magnetic stimulation coil for the rat.” Neuromodulation: Technology at the Neural Interface 19(5): 459–468.

68. Peterchev, A. V., et al. (2012). “Fundamentals of transcranial electric and magnetic stimulation dose: definition, selection, and reporting practices.” Brain stimulation 5(4): 435–453.

69. Poh, E. Z., et al. (2018). “Online LI-rTMS during a Visual Learning Task: Differential Impacts on Visual Circuit and Behavioral Plasticity in Adult Ephrin-A2A5–/–Mice.” Eneuro 5(1).

70. Rodger, J., et al. (2012). “Transcranial pulsed magnetic field stimulation facilitates reorganization of abnormal neural circuits and corrects behavioral deficits without disrupting normal connectivity.” The FASEB Journal 26(4): 1593–1606.

71. Rodger, J. and R. M. Sherrard (2015). “Optimising repetitive transcranial magnetic stimulation for neural circuit repair following traumatic brain injury.” Neural regeneration research 10(3): 357.

72. Rusjan, P. M., et al. (2010). Optimal transcranial magnetic stimulation coil placement for targeting the dorsolateral prefrontal cortex using novel magnetic resonance image-guided neuronavigation, Wiley Online Library.

73. Salvador, R. and P. Miranda (2009). Transcranial magnetic stimulation of small animals: a modeling study of the influence of coil geometry, size and orientation. 2009 annual international conference of the IEEE engineering in medicine and biology society, IEEE.

74. Saturnino, G. B., et al. (2019). “SimNIBS 2.1: a comprehensive pipeline for individualized electric field modelling for transcranial brain stimulation.” Brain and human body modeling: computational human modeling at EMBC 2018: 3–25.

75. Seewoo, B. J., et al. (2018). “Resting-state fMRI study of brain activation using low-intensity repetitive transcranial magnetic stimulation in rats.” Scientific reports 8(1): 1–13.

76. Shirinpour, S., et al. (2021). “Multi-scale modeling toolbox for single neuron and subcellular activity under Transcranial Magnetic Stimulation.” Brain stimulation 14(6): 1470–1482.

77. Siebner, H. R., et al. (2022). “Transcranial magnetic stimulation of the brain: What is stimulated?–a consensus and critical position paper.” Clinical Neurophysiology.

78. Stekic, A., et al. (2023). “Intermittent theta burst stimulation ameliorates cognitive deficit and attenuates neuroinflammation via PI3K/Akt/mTOR signaling pathway in Alzheimer’s-like disease model.” The alzheimer’s disease challenge, volume II 16648714: 44.

79. Suarez, L. E., et al. (2022). “A connectomics-based taxonomy of mammals.” Elife 11: e78635.

80. Suárez, R., et al. (2018). “A pan-mammalian map of interhemispheric brain connections predates the evolution of the corpus callosum.” Proceedings of the National Academy of Sciences 115(38): 9622–9627.

81. Tang, A., et al. (2017). “Repetitive transcranial magnetic stimulation of the brain: mechanisms from animal and experimental models.” The Neuroscientist 23(1): 82–94.

82. Tang, A. D., et al. (2021). “Subthreshold repetitive transcranial magnetic stimulation drives structural synaptic plasticity in the young and aged motor cortex.” Brain stimulation 14(6): 1498–1507.

83. Tang, A. D., et al. (2016). “Construction and evaluation of rodent-specific rTMS coils.” Frontiers in neural circuits 10: 47.

84. Thielscher, A., et al. (2011). “Impact of the gyral geometry on the electric field induced by transcranial magnetic stimulation.” Neuroimage 54(1): 234–243.

85. Wagner, T. A., et al. (2004). “Three-dimensional head model simulation of transcranial magnetic stimulation.” IEEE Transactions on Biomedical Engineering 51(9): 1586–1598.

86. Wang, B., et al. (2024). “Quasistatic approximation in neuromodulation.” arXiv preprint arXiv:2402.00486.

87. Weissman, J., et al. (1992). “Magnetic brain stimulation and brain size: relevance to animal studies.” Electroencephalography and Clinical Neurophysiology/Evoked Potentials Section 85(3): 215–219.

88. Yates, N. J., et al. (2014). “The role of ephrin-A2 and ephrin-A5 in sensorimotor control and gating.” Behavioural brain research 275: 225–233.

89. Yushkevich, P. A., et al. (2016). ITK-SNAP: An interactive tool for semi-automatic segmentation of multi-modality biomedical images. 2016 38th Annual International Conference of the IEEE Engineering in Medicine and Biology Society (EMBC), IEEE.

90. Zimdahl, J. W., et al. (2021). “Excitatory repetitive transcranial magnetic stimulation over prefrontal cortex in a Guinea pig model ameliorates tinnitus.” Frontiers in Neuroscience 15: 693935.

