## Supplementary Material for "Translational modelling of low and medium intensity transcranial magnetic stimulation from rodents to humans"

The following is the supplementary material to this article:

### S.1: Additional FEM methodology

#### S.1.1: Coil placement and general E-field distributions

All rodent coils were placed approximately 0.5 mm above the scalp (to consider the distance created by coil casings) over the left hemisphere with the outer edge of the coil positioned 1 mm lateral to the sagittal suture. The HI-TMS Cool-40 Rat coil was placed 3.5 mm over the scalp at a 30-degree angle to accommodate the casing and curvature of the coil, the outer edge of the coil was positioned with the midline between the inner and outer diameter over the midline of the left hemisphere. The HI-TMS D25 Alpha BI coil was placed approximately 3.5 mm above the scalp with the focal point of the figure of eight over the midline of the left hemisphere. All human coils were placed 10 mm above the scalp in consideration of the casing and hair with the MC-B70 figure of eight coil focal point placed over the DLPFC and the S90 Standard placed so the midline between inner and outer diameter was positioned over the DLPFC. Current in all coil models in this study was applied in a direction to induce E-fields that ran anterior to posterior through the left hemisphere (and DLPFC in human models).

Supplementary Figure 1 shows the E-field distribution of all rodent and human coils used in this study when positioned over the centre of spheres to better portray surface distribution; stimulation parameters were kept constant between models and heat maps were unscaled so that E-field distributions could be clearly displayed over a flat surface. The mouse, rat, and human spheres were an average diameter of the respective brain x, y, and z dimensions from Table 2.3 and were 10.4 mm, 18.14 mm, and 156.47 mm respectively; all spheres had isotropic conductivity. Figure of eight coils were constructed by modelling two homogenised multi-turn coils with current running in opposite directions with half the voltage applied to each coil.

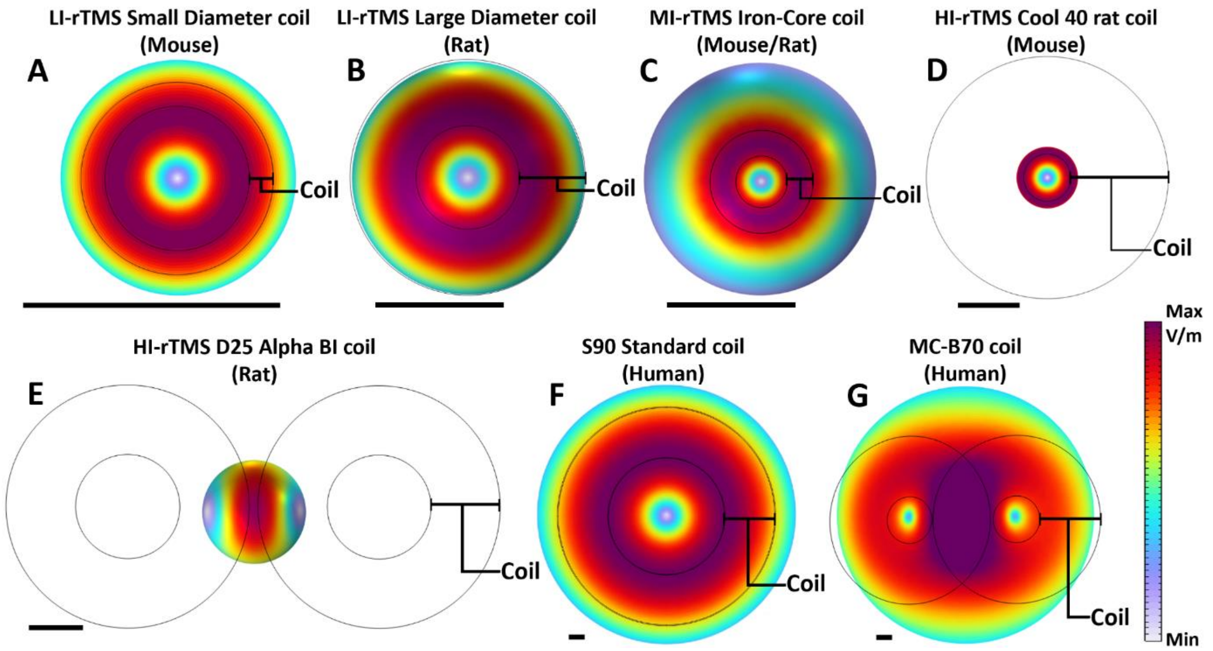


**Supplementary Figure 1.2:** E-fields of each coil type on species specific spherical model counterpart by all coils in this study. Spheres have a diameter the same as the length of the corresponding species brain. (A) LI-TMS mouse coil and MI-TMS mouse coil, (B) LI-TMS rat coil, (C) Rat iron-core coil, (D) 25 mm MagStim Rapid², © MagVenture MC-B70, (F) Circular human coil. Black bar = 10 mm. The MI-TMS Air-Core coil was omitted due to it having an identical E-field distribution to the MI-TMS Iron-core. Black bar in lower left of all images = 10 mm.

#### S.1.2: Additional finite element method equation, solver, and convergence details

The following set of equations was used in COMSOL for solving E (E-field V/m) and B (B-field T) in the frequency domain for all models used to calculate translational modelling results:

$$\nabla \times \mathbf{H} = \mathbf{J}$$

$$\mathbf{B} = \nabla \times \mathbf{A}$$

$$\mathbf{J} = \sigma\mathbf{E} + \mathrm{j}\omega\mathbf{D}$$

$$\nabla \cdot \mathbf{J} = \mathbf{0}$$

$$\mathbf{E} = -\nabla \mathrm{V} - \mathrm{j}\omega\mathbf{A}$$

For the constitutive relation B-H the relative permeability model was used to calculate **B** using:

$$\mathbf{B} = \mu_0\mu_\mathrm{r}\mathbf{H}$$

With $\mu_\mathrm{r}$ denoting relative permeability as a material property.

For the constitute relation Jc-E the electrical conductivity model was used to calculate $\mathbf{J}_\mathbf{C}$ using:

$$\mathbf{J}_\mathbf{C} = \sigma\mathbf{E}$$

For the constitutive relation D-E the dielectric model was used to calculate **D** using:

$$\mathbf{D} = \varepsilon_0\varepsilon_\mathrm{r}\mathbf{E}$$

With $\varepsilon_\mathrm{r}$ denoting relative permittivity.

The solver included a Coil Geometry Analysis as a preprocessing step to computer the current flow of a coil in 3D, followed by a stationary solver (MUMPS), the geometry shape function was Quadratic Lagrange, and the iterative solver was FGMRES.

The convergence plots generated by COMSOL for each of the 9 coils (LI-rTMS mouse and rat coils, MI-rTMS iron-core for mouse and rat coils, MI-rTMS air-core for mouse, HI-rTMS mouse and rat coils, the S90 standard coil, and the MC70-B coil) along with all translated modelling setups (S90 replicating rodent LI-rTMS and MI-rTMS) all returned a linear error result of 0.001 or below Convergence was tested by testing results of models with progressively larger element size and it was ensured that convergence of results was achieved prior to element sizes more than twice the final model sizes. In general, each model used element sizes as small as possible and element numbers for each model are given in the Results section of this paper.TMS E-fields as calculated by COMSOL are derived using Faraday's Law (Makarov, Yanamadala et al. 2015) with the magnetic vector potential (**A**):

$$\mathbf{E} = -\frac{\partial \mathbf{A}}{\partial \mathbf{t}} - \nabla\varphi$$

The tissue current is calculated using the addition of displacement current ($\frac{\partial A}{\partial t}$) and conduction current (σE):

$$\mathbf{J} = \sigma\mathbf{E} + \frac{\partial \mathbf{E}}{\partial \mathbf{t}}$$

#### S.2: Coil stimulation rise time (time domain) to frequency (frequency domain) conversions

The rise time of a triangle pulse is calculated from $T_o$ to $V_{peak}$ where V=0 at $T_o$. The rise time of a continuous sinusoidal wave in the frequency domain is nonlinear and results are taken with continuous stimulation assumed (e.g. including effects of hysteresis, without an initial magnetisation delay for iron-core coils).

The peak dI/dt (A/µs) with $dI/dt|_{Max}=V_c/L$ dI/dt is derived from the timeframe of pulse where the coil voltage changes most rapidly an amount equal to the maximum coil voltage. Frequency domain values were derived from time domain and real-life parameters using multiple data points that best approximated the same results at the same frequency for the same coil using the same parameters. This resulted in the following calculation:

$$\boldsymbol{f} = \frac{\mathbf{1}}{(\mathbf{RT}*\mathbf{0.6}*\mathbf{4})} \approx \frac{\mathbf{1}}{(\boldsymbol{RT}*\boldsymbol{\pi}*\mathbf{0.8})}$$

#### S.3: Air-core vs Iron-core rodent comparison

An axisymmetric FEM model was produced in COMSOL of both the iron-core and air-core MI-TMS rodent coils from this study (stimulated using the same parameters from this study). The spherical counterpart of the rat head model from Figure S.1 was used with tissue layers (representing the scalp, skull, CSF, GM, and WM). The sphere was placed 1 mm underneath the base of the MI-TMS coils and used to investigate the E-field and B-field distributions in both the frequency domain and time domain (Section 3.2). The time domain is a computationally demanding transient analysis that considers nonlinear magnetic properties of materials while models in the frequency domain assume a continuous sinusoidal coil excitation and don't typically include nonlinear magnetic properties. The axisymmetric model was constructed for time domain studies due to being less computationally demanding. Frequency and time domain studies were conducted using an air-core, an iron-core and an iron-core with BH-curve material properties calculated to approximate nonlinear magnetic excitation behaviour in the frequency domain (see Section S.3.1 below). These studies were threefold, to identify the accuracy of frequency domain studies compared to time domain studies, to identify how iron-core and air-core coils respond to pulse excitations across a range of rise times (or frequencies in the frequency domain), and to validate whether the Effective Nonlinear Magnetic Curves Calculator could produce an iron material for use in the frequency domain with more accurate behaviour.

#### S.3.1: The Effective Nonlinear Magnetic Curves Calculator

The default magnetisation model available in COMSOL for calculating the B-field in the frequency and time domain is the relative permeability model:

$$\mathbf{B} = \mu_0 \mu_\mathrm{r} \mathbf{H}$$

Where $\mathbf{B}$ is the magnetic flux density, $\mu_0$ is the magnetic constant, $\mu_\mathrm{r}$ is the relative permeability taken from the material properties, and $\mathbf{H}$ is the magnetic field strength. An alternative magnetisation model used in COMSOL based on a materials BH-curve (derived from real world testing) is the BH Curve model, which is only available in the time domain and given as:

$$\mathbf{B} = f(\|\mathbf{H}\|)\frac{\mathbf{H}}{\|\mathbf{H}\|}$$

This model describes the nonlinear behaviour of ferromagnetic material more accurately than the relative permeability model, taking into account real world data on specific materials. The Effective Nonlinear Magnetic Curves Calculator application is a companion to the Effective

Nonlinear Constitutive Relations functionality within the magnetic-based interface of the AC/DC Module in COMSOL and can be used to approximate the behaviour of a nonlinear magnetic material in the frequency domain. The Effective BH Curve model is available in the frequency domain and follows the same equation the BH Curve model. Figure S3.1 below shows the response under harmonic excitation of a nonlinear material (iron) under the most realistic time dependant BH Curve model (blue), the frequency domain permeability model (green), and the frequency domain effective BH curve model using an iron material produced by the Effective Nonlinear Magnetic Curve Calculator (effective iron material). This effective iron material was input as the material for cores in all iron-core coil models in this study using the Effective BH Curve model. The effective iron material was also used in the time vs frequency domain and iron vs air core studies seen below in section s.3.2 and replicated the time domain studies more closely (likely being more accurate).

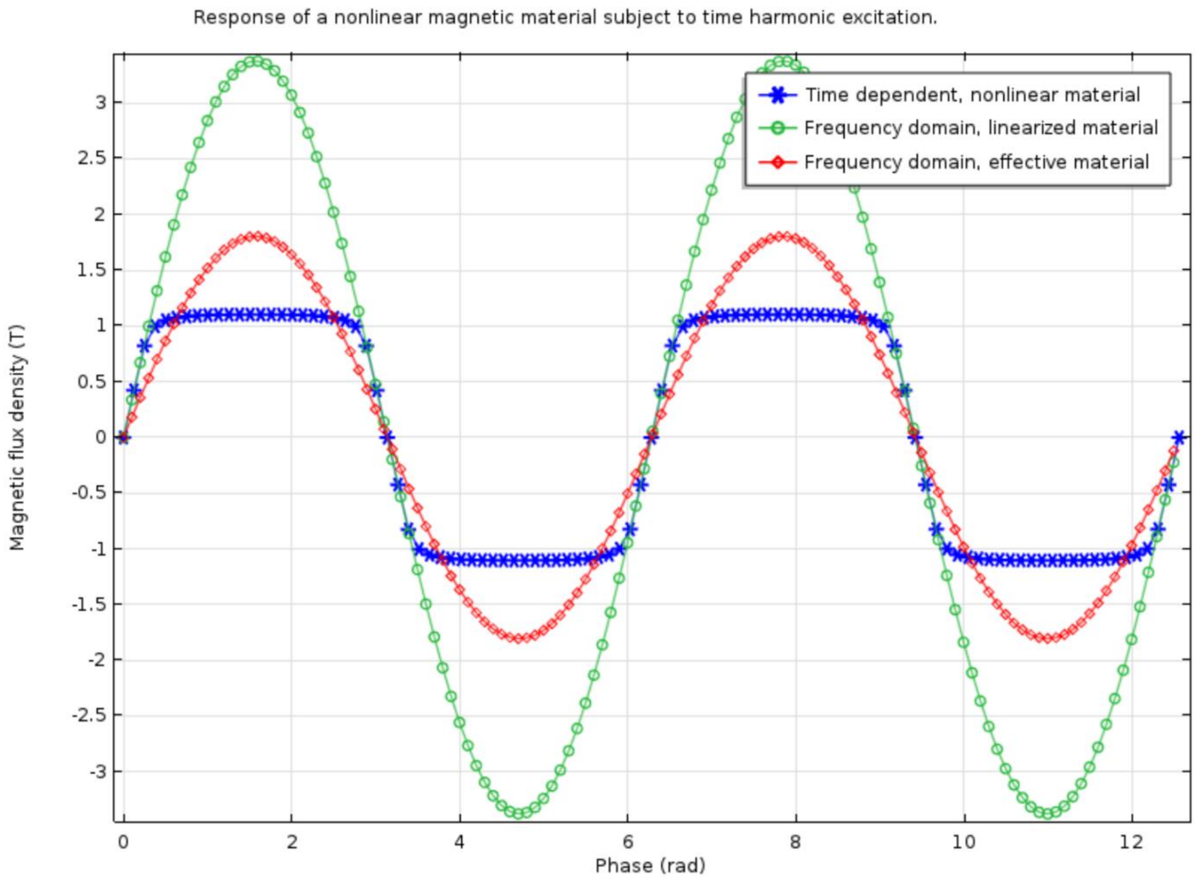


**Supplementary Figure 3.1:** Magnetic flux density (Tesla) response of a nonlinear magnetic material (iron) under harmonic excitation in the frequency domain (green, using linearised material properties), time domain (blue, including nonlinear material response), and in the frequency domain using a material with an effective BH curve calculated by the effective nonlinear magnetic curves calculator used to approximate the nonlinear behaviour of iron in the frequency domain.

#### S.3.2: Linear vs nonlinear material coil cores in the time and frequency domain

Figure S3.2.1 describes the B-field values produced at the GM surface directly under the centre of the TMS air-core and E-field values produced at the GM surface directly under the centre line between the inner and outer diameter of the base of the coil for the air-core, iron-core, and effective iron-core MI-TMS rodent coils, from this study, while being stimulated by an 100 V voltage source in the frequency domain by a range of frequencies between 0 kHz and 5.5 kHz. The air-core coil produced a weaker B-field than both iron-core coils under 0.5 kHz, a very similar B-field between 0.5 kHz and 1.5 kHz and a marginally stronger B-field over 1.5 kHz. The effective iron-core coil produced a weaker B-field than the regular iron-core coil at lower frequencies (>0.5 kHz) and due to its manner of calculation this likely a more realistic representation of the real thing. With lower frequencies (<0.75 kHz) the iron-core coils produce a stronger E-field, however, at frequencies over 0.75 kHz the air core produced an increasingly stronger E-field than both iron core counterparts. This was unintuitive as iron-core coils are generally assumed to produce stronger E-field values due to the stronger B-field values produced. The reasons for comparatively weaker E-fields in iron-core rodent coil models with coil excitation of frequencies over 0.75 kHz are not fully known, however, they are perhaps due to initial magnetisation delays or the iron-core saturation level. Stronger iron-core B-field values at frequencies below 0.75 was expected due to the higher capacity to concentrate magnetic flux lines.

Figure S3.2.2 describes the B-field and E-field values, measured from the same points as described for Figure S3.2.1 above, for the air-core, iron-core, and effective iron-core MI-TMS rodent coils, from this study, while being stimulated by a 100 V triangular pulse in the time domain by a range of pulse rise times between 0 µs and 4700 µs. The time domain models were reflective of the results in the frequency domain with a stronger B-field for both iron-core coils than the air-core coil for rise times over 250 µs, while again, below 250 µs air-core and iron-core coils induced very similar B-field values. Very similar E-field distributions across the air-core and iron-core coils were seen in the time domain when compared to the frequency domain with the air-core coil producing a stronger E-field at rise times below 250 µs but the air-core producing stronger E-fields at rise times above 250 µs.

These results tend to suggest that air-core rodent coils of this size produce stronger E-field values than iron-core coils of the same size when pulses with rise times below approximately 250 µs are applied.

Interestingly, due to hysteresis the iron-core coils produced a far weaker E-field during the fall time when compared to the air-core coils. It should therefore be assumed that the strongest

E-field is produced in an iron-core coil during the rise time, however, this may present an issue when biphasic pulses are used as a means of creating a stronger E-field as necessarily the fall time and subsequent reversing of polarity is relied upon to produce the strongest E-field across the pulse. A way to avoid hysteresis would be to use a laminated iron core in place of a solid iron core or to produce a coil with a ferrite core, which would be unaffected by hysteresis.

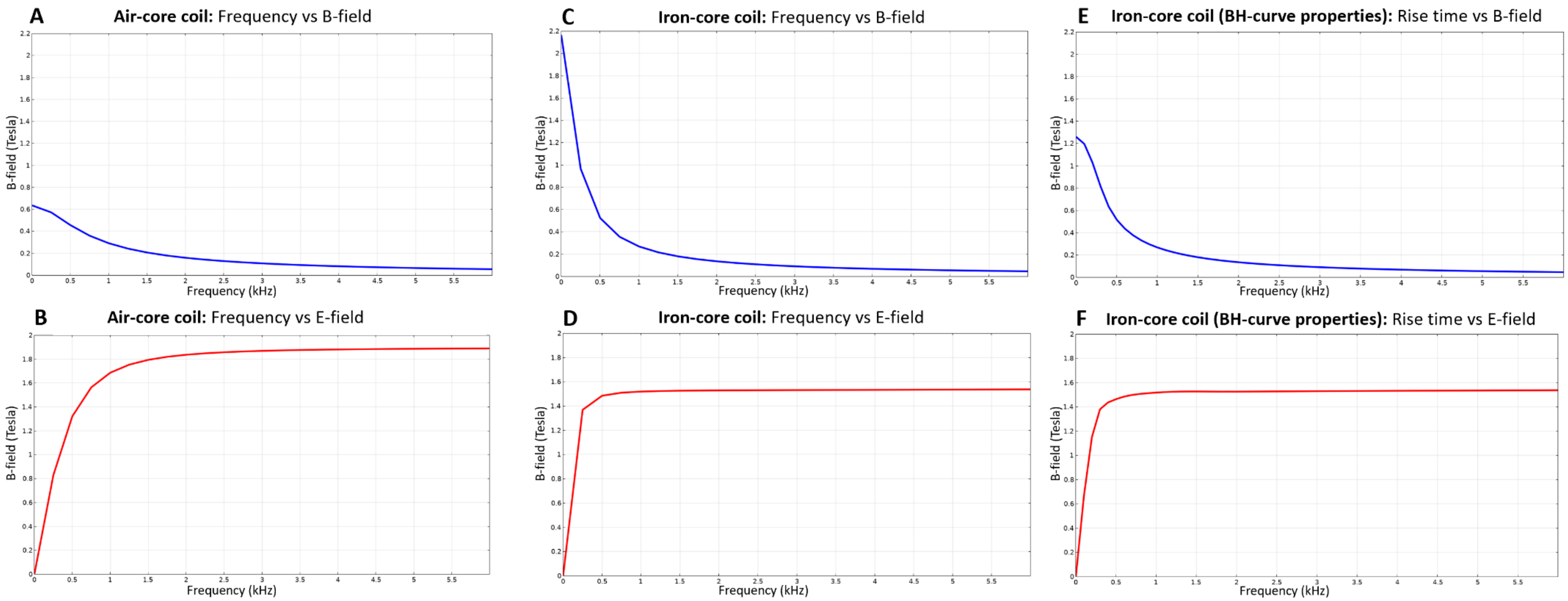


**Supplementary Figure 3.2.1:** Graphs showing the frequency vs B-field for the (A) air-core, (C) iron-core, and © effective iron-core coils and the frequency vs E-field for the (B) air-core, (D) iron-core, and (F) effective iron-core coils.

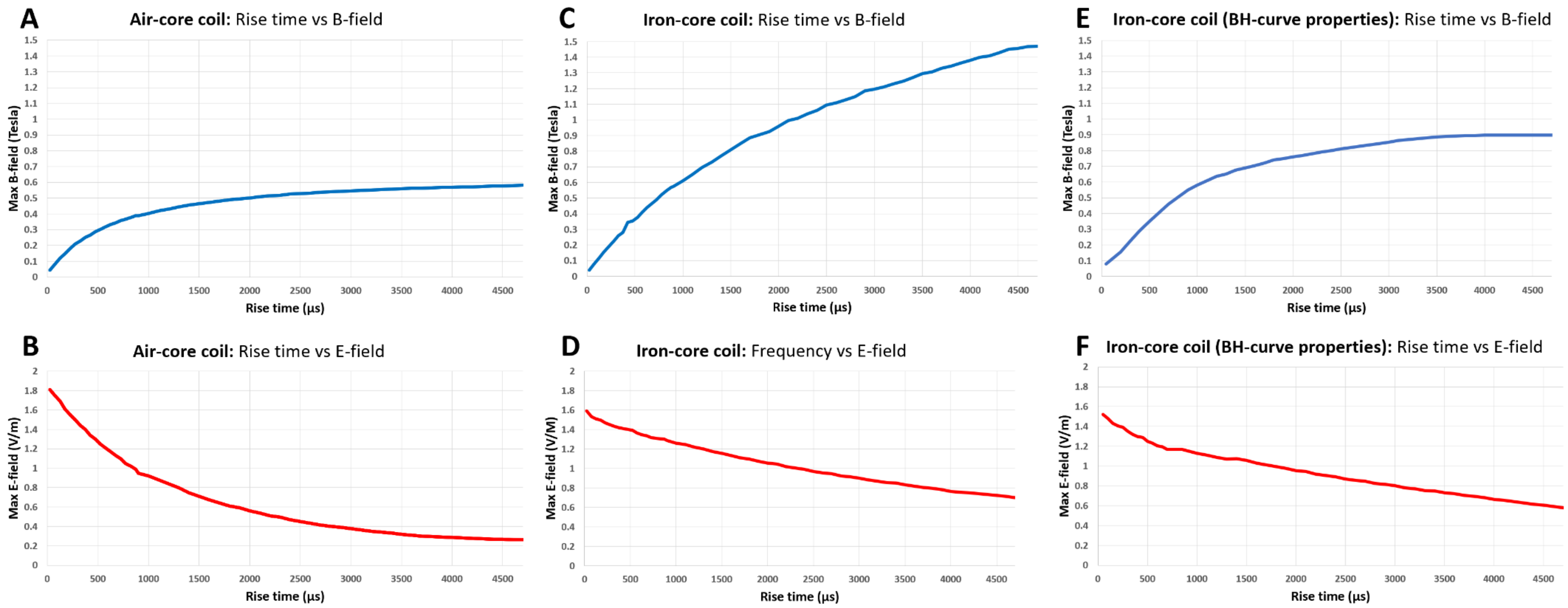


**Supplementary Figure 3.2.2:** Graphs showing the rise time vs B-field for the (A) air-core, (C) iron-core, a©(E) effective iron-core coils and the rise time vs E-field for the (B) air-core, (D) iron-core, and (F) effective iron-core coils.

#### S.3.3: Laboratory measurements of air-core vs iron-core rodent coils

A Hall Effect probe (Honeywell SS94A2D, NJ, USA) and a stereotaxic frame (for coil and probe positioning) was used to measure the magnetic field (B-field) produced both the MI-TMS air-core and MI-TMS iron-core rodent coils used in this study at 2 mm increments between 0 mm and 10 mm directly away from the base of the coil with coil centres positioned directly above the probe (xy =0 mm, z = 0-10 mm). Both coils were stimulated using a triangle pulse with a rise time of 200 µs produced by a wave generator (Keysight 33500B Series, USA) set to three different output values (Vpp – 1.5 V, 3 V and 5 V), which were input into and amplified by a power supply (Kepco, BOP 100-4M, NY, USA). These data were collected to ascertain both the difference in B-field strength between the air-core and iron-core coils as well as the B-field strength over distance, the later measurements being collected in light of the respective iron-core coil FEM frequency domain model showing "tighter" flux lines in comparison to the air-core coil. There were locations across space where the iron-core coil model produced a stronger B-field in comparison to the air-core coil while other locations across space the air-core's B-field was comparatively stronger (the later occurring further from the base of the coil). This effect could be on account of the iron-core coil producing less leakage flux and fringing with the iron-core presenting a more magnetically conductive pathway where flux lines are gathered closer leading to a larger B-field gradient moving away from the base of the coil. However, the measurements showed that while the B-field produced by the iron-core coil was clearly stronger at close distances (see Figure S3.4) the difference in magnitude was reduced over distance to approximately the same values at 10 mm distance from the base of the coil.

The wave generator was set to the same values for both coils, however, the air-core drew less voltage, while adjustments could be made at the two lower voltages the maximum output for the waveform generator was chosen for the highest voltage and manual adjustments beyond this, via the power supply, would cause the rodent coils to produce excessive heat and be damaged beyond use.

It should be noted that the iron-core coil used for the measurements in Figure S

3.3 had a resistance of 21.8 ohms and a measured inductance (using a Micron Q1150B digital LC meter) of 11.22 mH, while the air-core coil measured 24.8 ohms and 1.86 mH, the COMSOL model counterparts measured 6.26 mH and 1.54 mH respectively. Deviations in the process of manual iron-core construction of rodent coils can alter the inductance value as can exposure do mechanical damage and high operating temperatures with another older well used iron-core coil measuring 24.5 ohms and 2 mH.

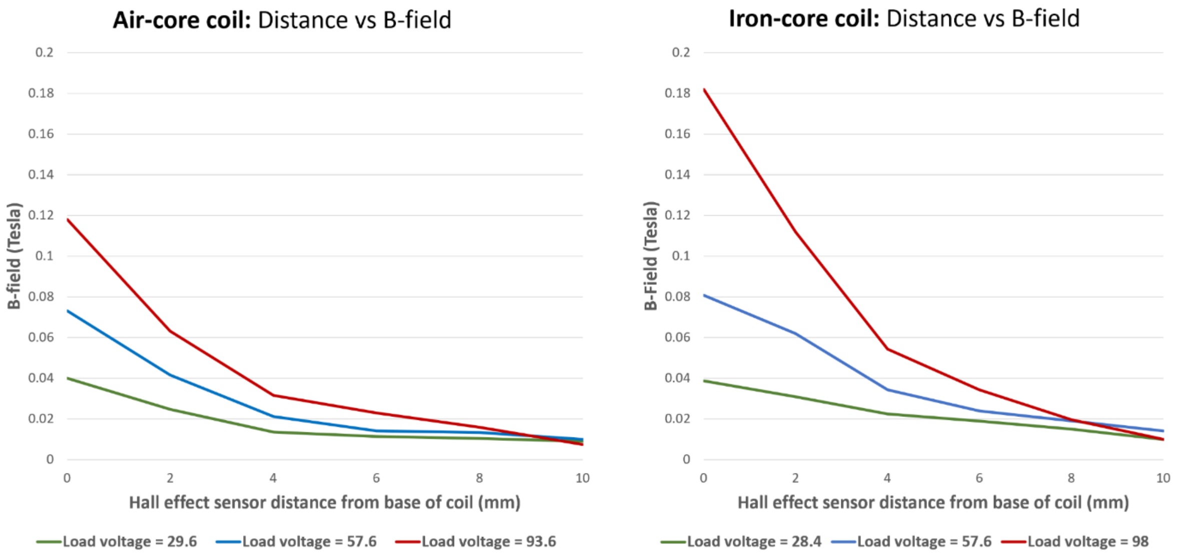


**Supplementary Figure 3.3:** Air-core (left) and iron-core (right) B-field magnitudes over distance with coil centres positioned directly above the probe. Three voltage values were used for coil stimulation and are listed in the legend below each graph, the maximum output stimulating the coils differed with the iron-core drawing a higher coil voltage.

### S.4: Electrical circuit physics module details

To improve the accuracy of FEM modelling results for the rodent TMS the circuit diagram used in the related experiments was modelled using COMSOL's Electric Circuit physics interface. The circuit diagram included the source impedance and resistance present in the Kepco, BOP 100-4M power supply (which ultimately had a negligible effect on results) and the coil impedance (see Figure S4.1). Furthermore, the voltage source for time domain simulations included a pulse, which is described in part B of Figure S4.1.

During these studies modelling work it was identified that the purely inductive load of the rodent coil TMS setup the power factor is approximately 0.5. The reactive power (wasted power) being reflected away from the coil can be reduced and the active power (and E-field distribution), can be increased by adding a capacitor in parallel to the coil (see Figure S4.1 part **C**), which reduces the power factor and according to modelling results would increase the E-field (in our FEM models by 50% to 100%). Various capacitor values were tested in the time domain using the axisymmetric iron-core rodent coil model with a 100 V triangle pulse and 75 µs rise time (Figure S4.2 for E-field values and Figure S4.3 for B-field values) with B-field and E-field values being collected at the same locations as in Section S.3.2. E-field values were increased with the increasing parallel capacitance value between 0 and 5000 nF, the E-field continually decreased with capacitances over 5000 nF. However, as observed in Figure S4.2, adding the capacitor in parallel creates harmonic oscillations, these oscillations increased with the increasing capacitor. A capacitance value between 1000 nF and 2000 nF produced minimal oscillations. The air-core coil with the capacitor in parallel produced higher E-fields (approximately by 2-5%) than the iron-core. Adding a capacitor in parallel increases E-field values, likely without production of additional coil heat.

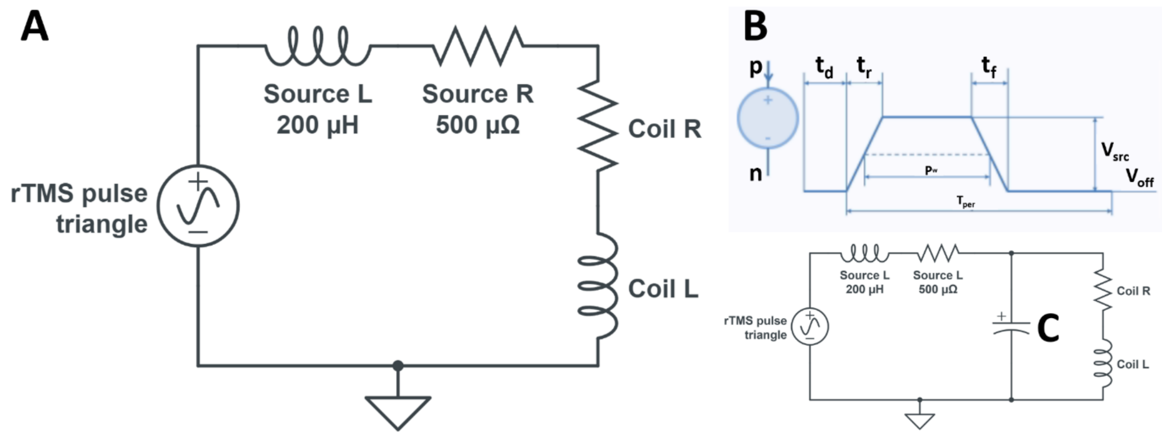


**Supplementary Figure S4.1:** Circuit (A) used to produce the triangle pulse (time domain) or continuous sine wave (frequency domain) for the rodent coil models in this study. The time domain pulse source (B) can be combined with more pulses to create a biphasic pulse. Adding a capacitor in parallel to the coil (C) improves the power factor and allows for the production of stronger B-fields and E-fields, however, this also results in harmonic oscillations in the circuit which are observed as multiple B-field an E-field pulses across time (see Figure S4.2).

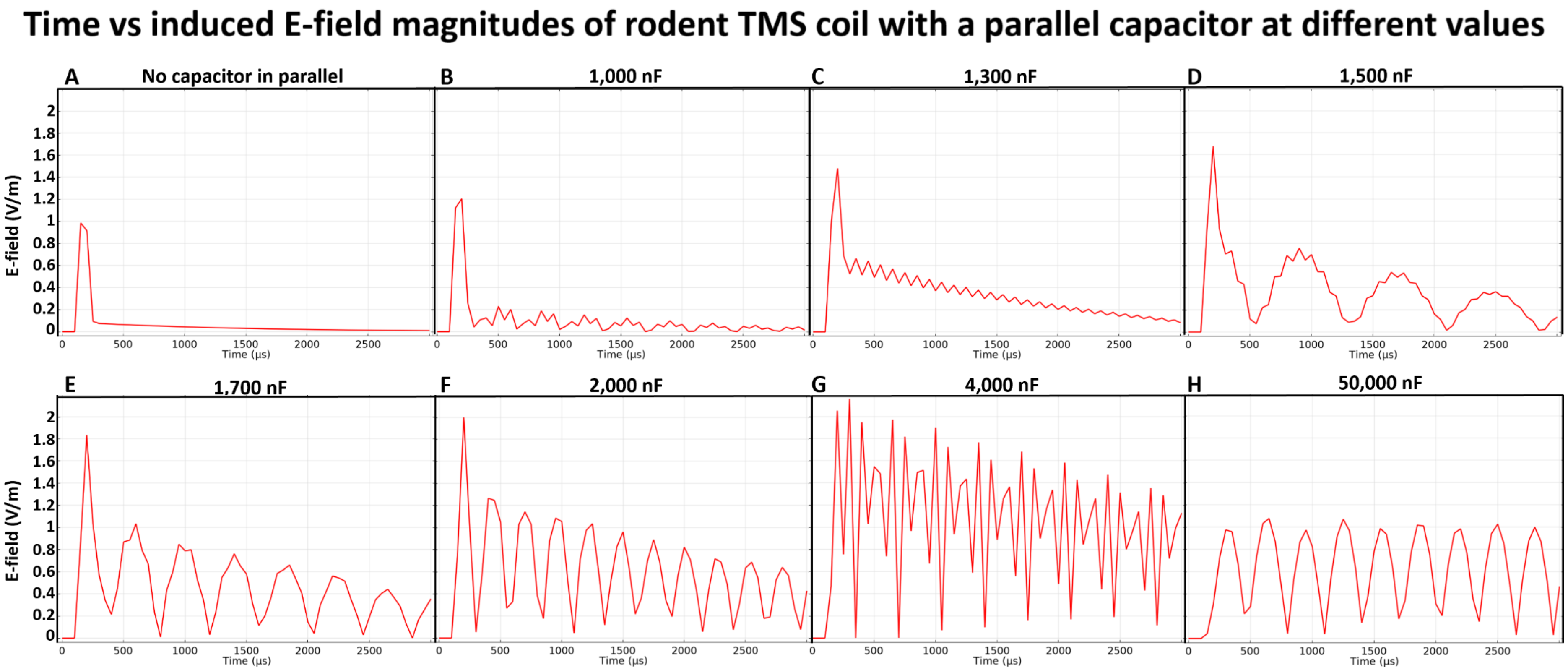


**Supplementary Figure 4.2:** E-field values observed at the GM of the spherical counterpart to the rat head model directly beneath the midline between the inner and outer diameters of the MI-TMS iron-core coil. Values are recorded for the (A) regular circuit (see Figure S4.1 part A) and with a capacitor in parallel with the coil with a value of (B) 1,000 nF, (C) 1,300 nF, (D) 1,500 nF, (E 1,700 nF, (F) 2,000 nF, (G) 4,000 nF, and (H) 50,000 nF.

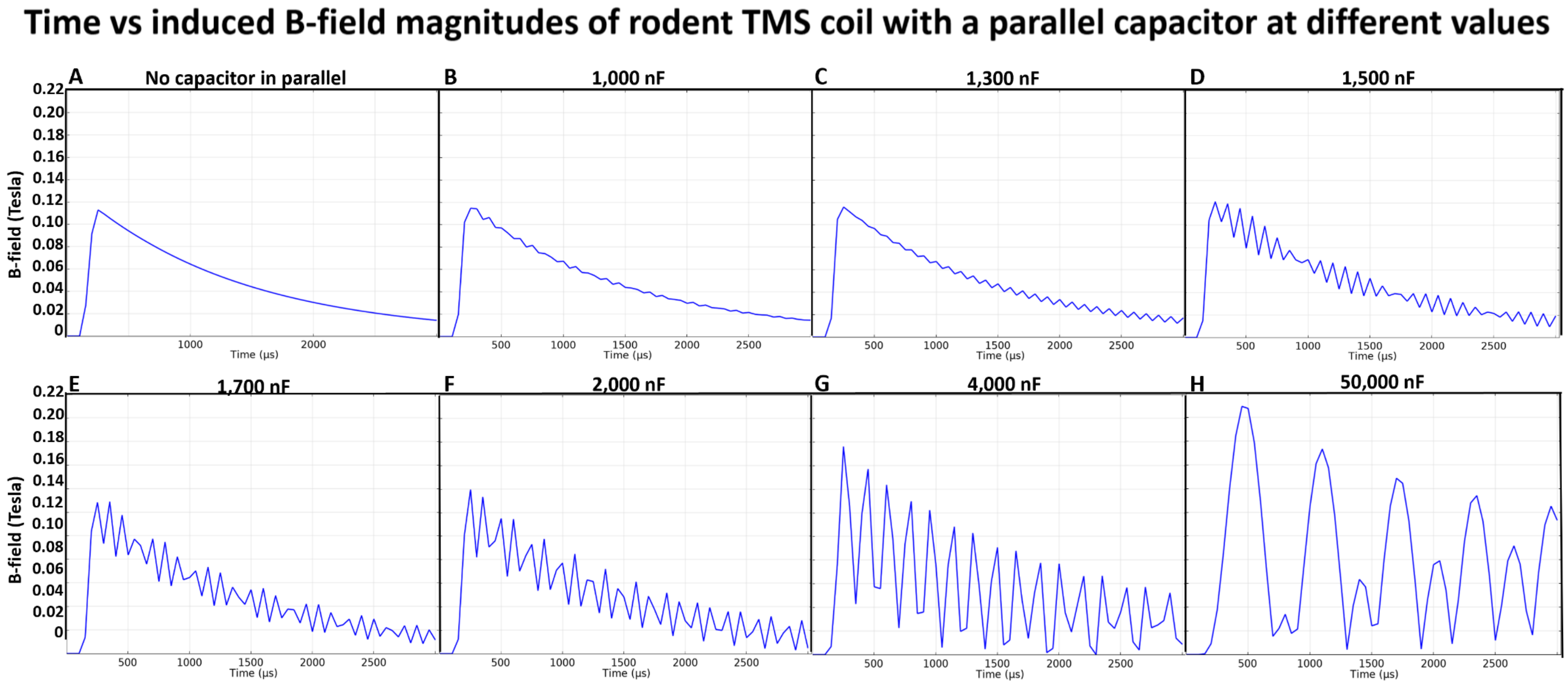


**Supplementary Figure 4.3:** B-field values observed at the GM of the spherical counterpart to the rat head model directly beneath the centre of the MI-TMS iron-core coil. Values are recorded for the (A) regular circuit (see Figure S4.1 part A) and with a capacitor in parallel with the coil with a value of (B) 1,000 nF, (C) 1,300 nF, (D) 1,500 nF, (E 1,700 nF, (F) 2,000 nF, (G) 4,000 nF, and (H) 50,000 nF.

### S.5: COMSOL vs SimNIBS E-field distributions

Included below are E-field and B-field peak values for the LI-rTMS and HI-rTMS mouse coils (Supplementary Figure 5.1) and for the LI-rTMS, MI-rTMS, and HI-rTMS rat coils (Supplementary Figure 5.2).

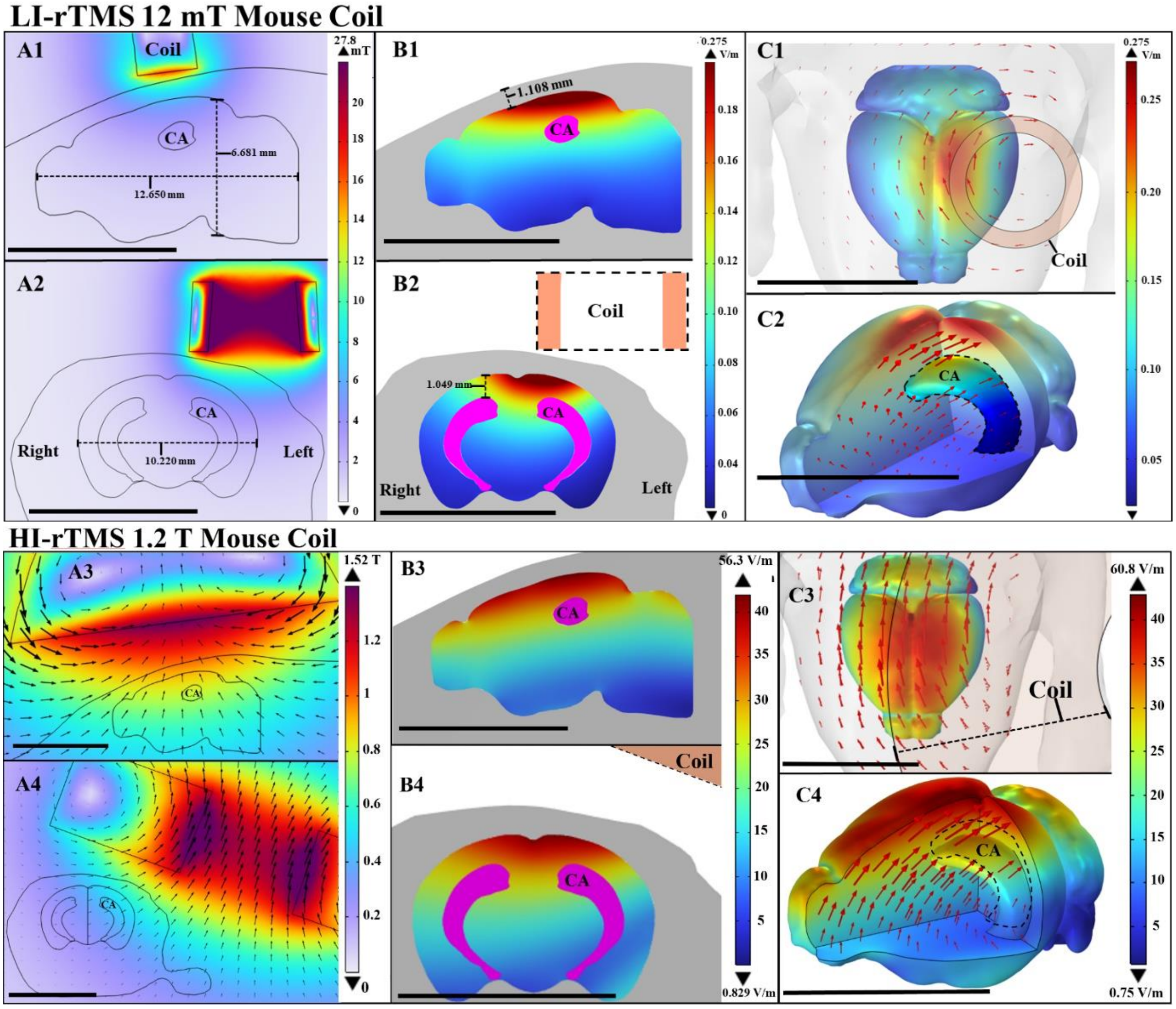


**Supplementary Figure 5.1:** Peak E-field and B-field heat maps shown for the 12 mT LI-TMS Small Diameter and 1.20 T HI-TMS Cool-40 rat (set to 20% MSO) coils. With A1 and A2 showing the 12 mT Mouse Coil-derived B-field for the sagittal and coronal planes respectively and B1 and B2 showing the same planes for the E-field maps. C1 and C2 are 3D views of the 12 mT Mouse coil E-field map. A3 and A4 are showing the 1.2 T Mouse Coil-derived B-field for the sagittal and coronal planes respectively and B3 and B4 showing the same planes for the E-field maps. C3 and C4 are 3D views of the 1.2 T Mouse coil E-field map. Scale bar in first two rows = 10 mm

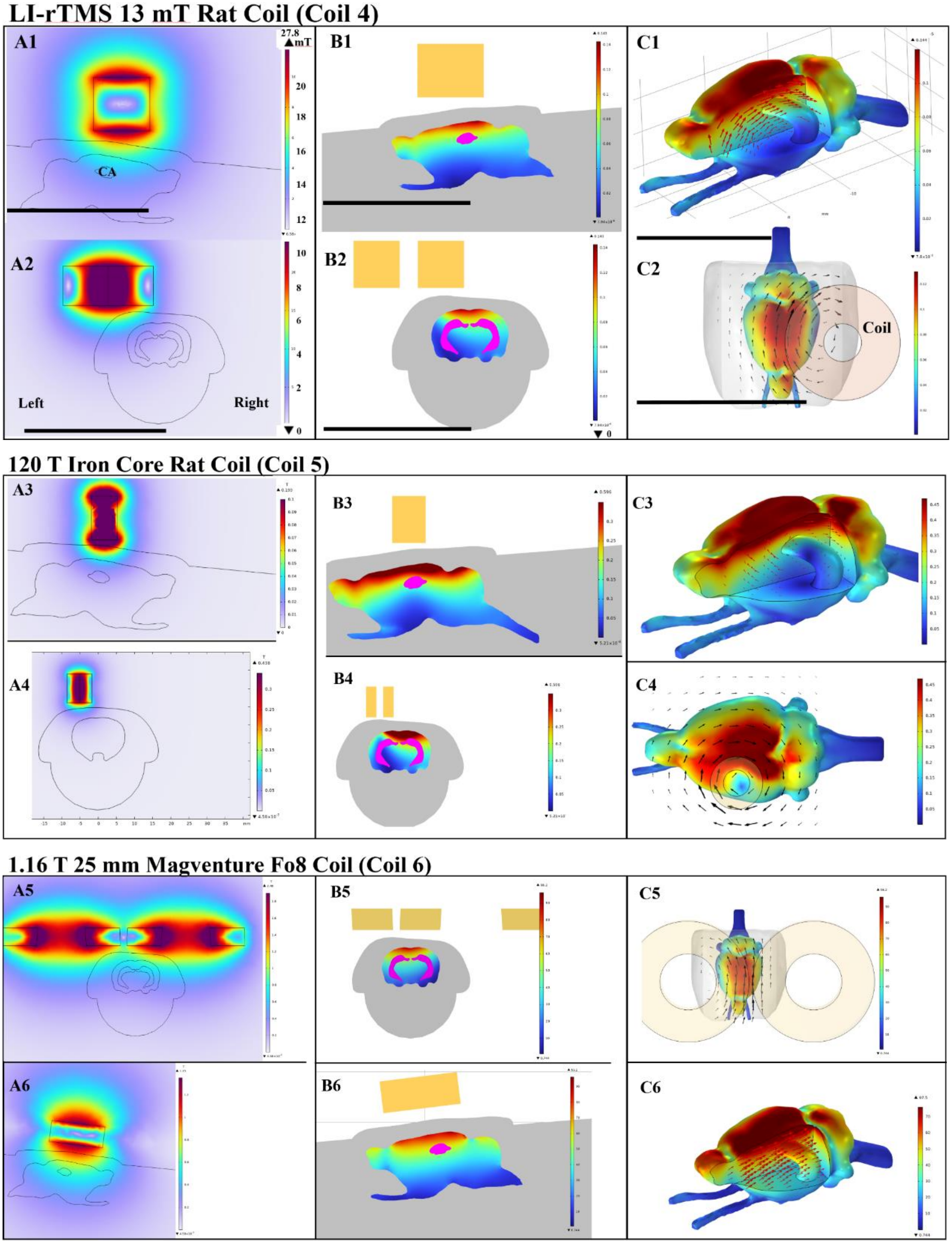


**Supplementary Figure 5.2:** Peak E-field and B-field heat maps shown for the 13 mT LI-TMS Large Diameter, 120 T Iron Core and 1.16 T 25 mm Magventure Fo8 (set to 33% MSO) coils. With A1 and A2 showing the 13 mT Rat Coil-derived B-field for the sagittal and coronal planes respectively and B1 and B2 showing the same planes for the E-field maps. C1 and C2 are 3D views of the 13 mT Rat coil E-field map. A3 and A4 are showing the 120 mT Iron-Core Rat Coil-derived B-field for the sagittal and coronal planes respectively and B3 and B4 showing the same planes for the E-field maps. C3 and C4 are 3D views of the 120 mT Iron-Core coil E-field map. A5 and A6 are showing the 1.16 T 25 mm Magventure Fo8 Coil-derived B-field for the sagittal and coronal planes respectively and B5 and B6 showing the same planes for the E-field maps. C5 and C6 are 3D views of the 1.16 T 25 mm Magventure Fo8 Coil E-field map. Scale bar in first two rows = 10 mm

#### S.6: COMSOL vs SimNIBS E-field distributions

SimNIBS (Version 4.0.0) was used to simulate TMS of the S90 Standard and MC-B70 coils at maximum stimulator output using the same MNI152 head model from this current study. SimNIBS models were run at 50% MSO for both coils and at each of the MSO % settings and distances used in section 3.5 and Figure 3.8 (See Supplementary Figure 2). Due to the method of deriving E-field values in SimNIBS, when the coil is placed at 133- and 177-mm additional distance the SimNIBS coil models introduced E-field values at zero value beyond the dimension of space covered by the coil file (see S6 and S7 in Supplementary Figure 2). As such using SimNIBS in its current form (Version 4.0.0) with the aim of calculating E-fields when coils are placed at distances, such as in this paper, will not describe the E-field in its entirety.

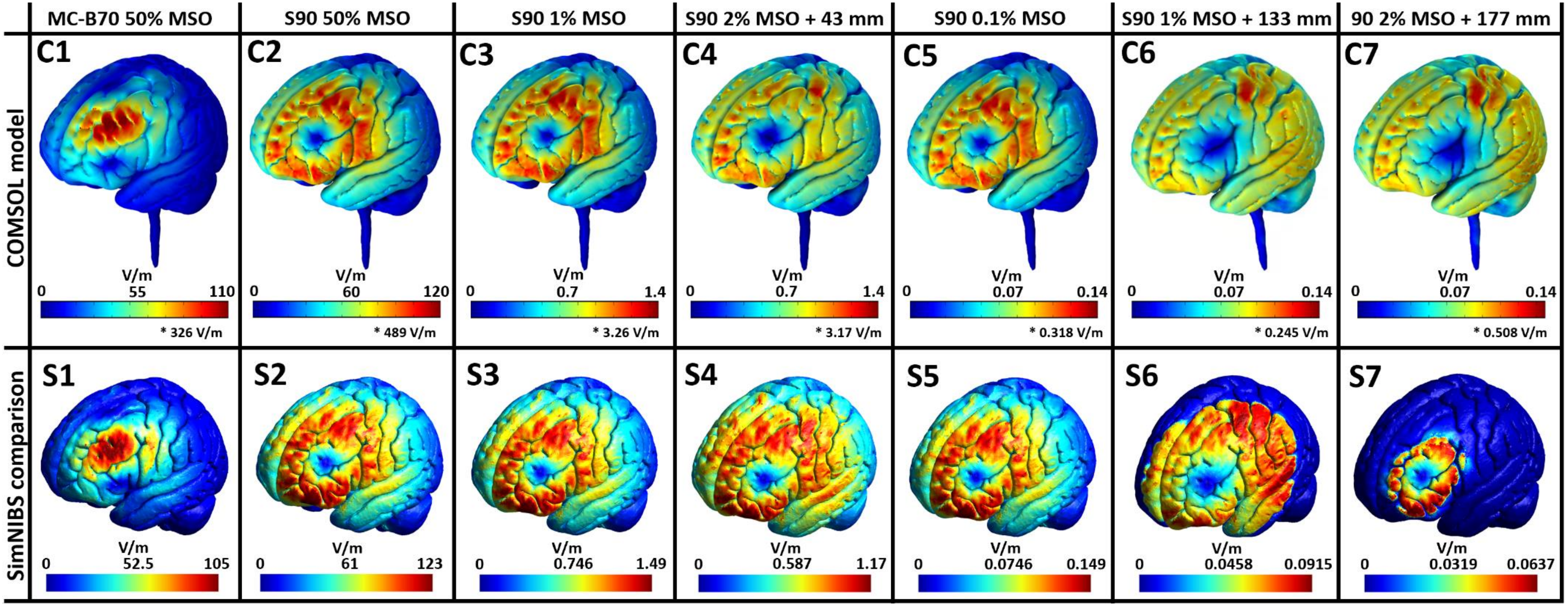


**Supplementary Figure 6.1:** TMS simulations of the MC-B70 coil induced E-field in the MNI152 human brain model from this study with the stimulator set to 50% MSO in both COMSOL (C1) and SimNIBS (S1). Simulations of the S90 Standard coil induced E-field were also produced in COMSOL with the stimulator set to 50% MSO in COMSOL (C2) and SimNIBS (S2). To compare E-fields induced in MI-TMS rodent studies the S90 was set to both 1% MSO and 2% MSO with the coil distance increased by 43 mm for both COMSOL (C3 and C4 respectively) and SimNIBS (S3 and S4 respectively). To compare E-fields induced in LI-TMS rodent studies the S90 was set to both 0.1% MSO, 1% MSO with the coil distance increased by 133 mm and 2% MSO with the coil placement increased by 177 mm for both COMSOL (C5, C6 and C7 respectively) and SimNIBS (S5, S6 and S7 respectively).

*COMSOL models include a small number (generally 3-5) miniscule (~1 $mm^3$) locations where peak E-field values jump significantly, likely due to a combination of sharp edges gathering charge and these locations being the smallest distance across a tissue of lower conductivity. These higher peak values are placed above COMSOL legend bars.

### S.7: Additional TMS coil modelling parameter details

**Table S7:** Parameters considered integral to electromagnetic FEM model accuracy for representing realistic TMS phenomena in the frequency domain using COMSOL.

| Parameter name | Importance | Issues | Recommendations |
|---|---|---|---|
| Frequency | Medium | Assumptions are built into frequency selection as the TMS pulse is interpreted as a continuous AC signal. | Calculated based on extrapolating from the fastest changing segment of the pulse |
| Physics interface choice | Medium | Some interfaces are missing mathematical detail. | Use the Magnetic and Electric Fields interface |
| Physics nodes choice | High | Equations involved vary between nodes changing what variables are included, what mathematical concepts are considered or omitted, computational time, result visualisation. | Use Ampere's Law and Current Conservation unless mesh resolution is extremely high or current losses are a focus |
| Mesh resolution | Medium | Accuracy of results are directly related to this factor. | Decrease the mesh size continually by half until results remain consistent for at least 2 solutions in a row. Resolution needs to be high between the coil and through the entire head model. |
| Mesh type | Medium | Changes accuracy and computational time | For complex meshes such as those involved in this study meshing outside of COMSOL can prevent issues. |
| Material properties (conductivity, relative permeability, and relative permittivity) | Mouse | Sources are minimal but preserved through the literature. The required rodent data is not available. | Use databases referenced in this study until rodent data becomes available. Air has a very small conductance. |
| Features of curved geometry | High | Sharp boundaries contribute to model result artifacts unless they're realistic. | Increase curvature where possible as much as is allowed. |
| Boundary conditions | Human | These are integral to how E-field results are calculated within volumes | This varies widely with the focus of the study and depends on if other types of physics are present in the model. |
| Outer boundary conditions | Medium | The magnetic field will interact with an outer boundary, it's important there are no unrealistic outcomes due to this factor. | An infinite boundary prevents the magnetic field reflecting and interfering with results. |
| Coil conditions | High | The coil can be stimulated by a range of methods in COMSOL. | This varies widely with the focus of the study. |
| Results | Medium | Results extracted from boundaries are more prone to artifacts. | If possible, note any differences between results central to a volume and those at the boundary. 2D results can change rapidly across the third axis so positioning is integral. |

### S.8: Construction of the mouse, rat, and human head models

A 3D FEM mouse model was created by combining the anatomical atlas of the adult male nude mouse "Digimouse" (Dogdas, Stout et al. 2007), with higher resolution MRI images of the P56 mouse atlas brain (Lein, Hawrylycz et al. 2007). Using the FMRIB software library (Jenkinson, Beckmann et al. 2012) Brain Extraction Tool (BET), the P56 brain was scaled and mapped to the original brain space and the "Digimouse" spinal cord was kept intact. The mouse brain model dimensions were in line with the average volume for an adult mouse (Hammelrath, Škokić et al. 2016)

The rat head model was derived from high resolution T2 weighted anatomical images using isotropic voxels of an ex-vivo head of a Sprague-Dawley rat (weighing 280g), collected using a 9.4T Bruker Biospec 94/30 small animal MRI machine (Figure S8.1). 3D surfaces for the scalp, skull, and GM were segmented out using ITK-SNAP (Yushkevich, Gao et al. 2016). The Waxholm space atlas of the Sprague Dawley rat brain was used to section out brain regions (Papp, Leergaard et al. 2014). The rat brain model dimensions were in line with the average volume for an adult Sprague Dawley rat (Goerzen, Fowler et al. 2020).

A 3D FEM human head model was produced using CerebrA (Manera, Dadar et al. 2020), an updated version of the 2009 symmetric MNI-ICBM152 (MNI152) coordinate system (Fonov, Evans et al. 2009, Fonov, Evans et al. 2011); Brain regions were section out using the MNI structural atlas (Collins, Holmes et al. 1995, Mazziotta, Toga et al. 2001) and the Harvard-Oxford cortical and subcortical structural atlases (Frazier, Chiu et al. 2005, Desikan, Ségonne et al. 2006, Makris, Goldstein et al. 2006, Goldstein, Seidman et al. 2007).

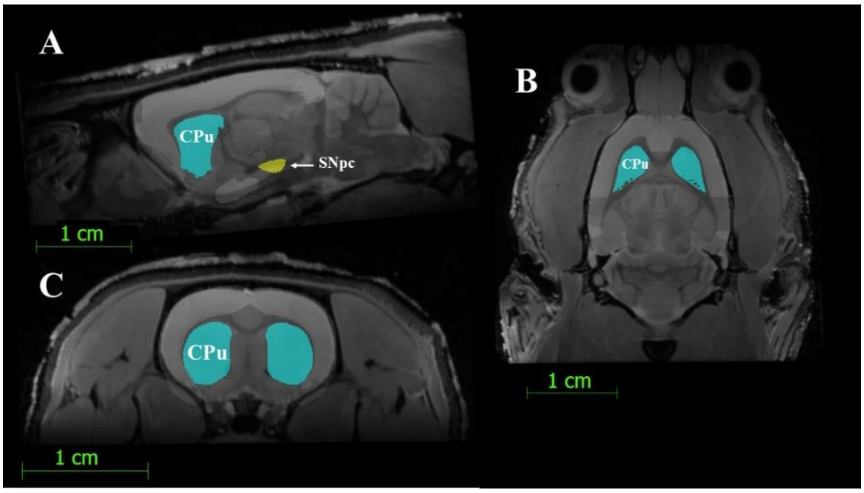


**Supplementary Figure S8.1:** T2 weighted MRI images captured for this study and used to develop the 3D rat head model. The sagittal (A), frontal (B) and transverse (C) planes are shown with the caudate putamen (CPu) and substantia nigra pars compacta (SNpc) brain regions shown highlighted for sectioning out as a separate 3D model.

#### S.9: Additional modelling limitations and artifacts

In the process of building FEM model head geometry smooth surfaces of tissues are approximated by polygons with edges and vortices. This approximation mixed with resolutions of source images, computational limitations (for 3D model meshing) and the assumptions made by the chosen physics interfaces results in unavoidable modelling artifacts. For example, the rat model was derived from a cube shaped MRI image and required a simple smooth geometry extension to accommodate missing body sections with cropped models impacting E-field distribution significantly (Mantell, Perera et al. 2023). Two noticeable artifacts in the models from this study are, firstly, how sharp angles in the geometry of dielectrics alter the E-field distribution and, secondly, how even thin layers of tissue with relatively large differences in dielectric properties can change the overall E-field distribution.

##### S.9.1: Sharp angles in FEM geometry alter the E-field distribution

Sharp angles in tissue geometry cause localised increases in E-field magnitude and gradient and these have been described in previous FEM modelling of TMS (Makarov, Yanamadala et al. 2015). These increases are proportional to the difference in conductivity and "sharpness" of the angle between the two tissues with the tissue constituting the convex aspect of the angle experienced the biggest change. Simplified FEM geometry causing unrealistic angles is therefore prone to artifacts, however, there are locations where this effect is likely present in real life. For example, in the rodent brain models where the most caudal location along the midline of the cortex meets the most rostral part of the midbrain an angled concave surface is formed causing an approximately 37.5% local increase in E-field magnitudes, which are not representative of the broader E-field distribution, but the geometry is similarly sharp in the biological rodent brain and therefore more likely to exist. Further experimentation is required to ascertain whether this is the case.

Similar to rodent counterparts, the human brain model featured a section of sharp geometrical surface of approximately 2 $mm^2$ where a significant jump in E-field magnitude was observed up to 213 V/m and 244.5 V/m for the MC-B70 and S90 models respectively. Furthermore, this location was within a sulcus at adjacent GM surfaces and 5 other locations presented similar results of approximately 150 V/m, ultimately these results were removed from Figures due to lack of clarity due to the high likelihood that these jumps in magnitude are due to modelling artifacts since the real human brain is devoid of sharp angles but the complexity of its curves leads to sharp angles in the lower resolution FEM counterpart (based on an aggregation of tetrahedra) due to lack of computational resources. In future, a supercomputer running the same human FEM brain model with a much higher resolution would remove geometric artifacts.

Values were taken from FEM models at three-dimensional 0.25 mm grid points from rodent models and 1 mm grid points from human models. This method reduces the effect of increased E-field values seen at sharp angles, where angles of artificially increased sharpness are created in 3D FEM models at borders between dielectrics (tissue layers) of the head models due to computationally constrained mesh resolutions. By taking values away from tissue borders at grid points these artificially increased E-field values are removed to an extent, otherwise the sharpest angle present in any brain region would likely hold an artificially increased E-field.

#### S.9.2: The inclusion of CSF conductivity improves accuracy of FEM models of TMS

The high conductivity of CSF allows the E-field to travel further along the brain surface and generate higher magnitudes at surface points than would otherwise be present. However, there is no standard CSF volume for a rodent FEM model and using individual MRI images for CSF volume estimation in rodent models can be misleading, especially when dealing with deceased or injured specimens.

This study and previous modelling efforts (Alekseichuk, Mantell et al. 2019), have included a assumed volume of CSF, however, high definition structural MRI imaging of rodents from a healthy and representative population can provide more detail on how to model the CSF for FEM purposes.

#### S.9.3: The difference in the depth and location of brain regions lead to issues in E-field replication across species

Figure S9.1 provides a visual representation of how cortical folding, differences in brain region location, and an increase in overall size of the brain all provide issues in reproducing E-field values in multiple brain regions across species.

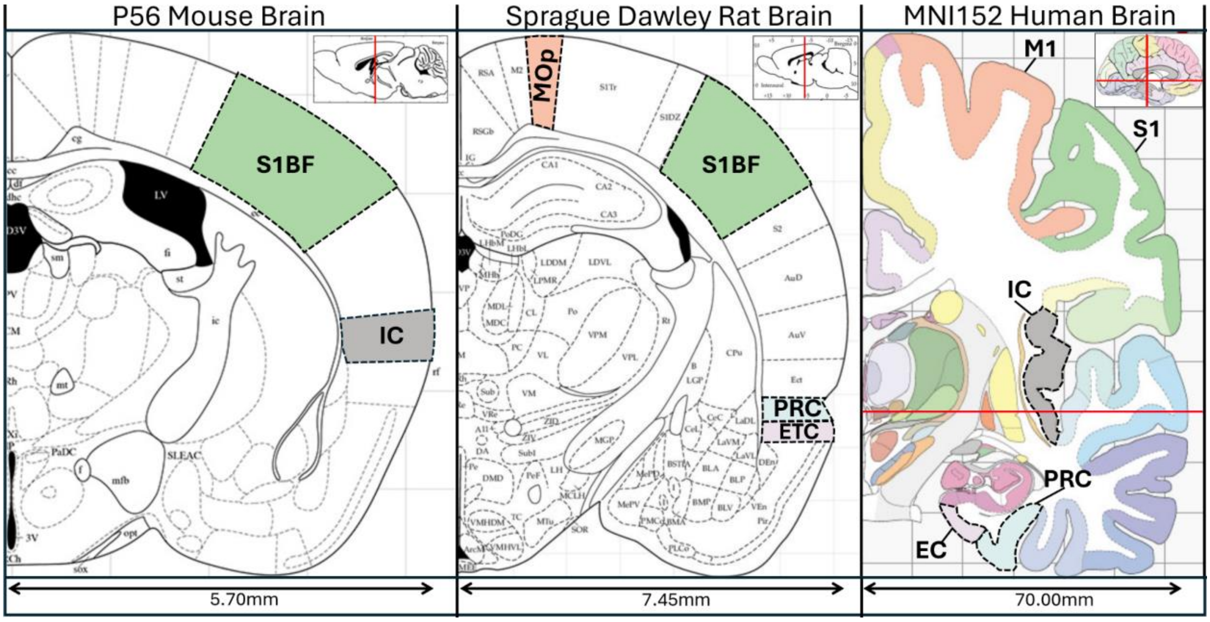


**Supplementary Figure S9.1.** Brain region atlas of comparative brain regions for a mouse, rat, and human. While some brain regions are located in anatomically similar position with respect to gross brain anatomy across species (e.g. somatosensory and motor cortexes), brain folding in primates has increased the depth of some brain regions (see insular cortex) while the extended cortex of primates means other regions are located more inferiorly and medially compared to rodent regions (see entorhinal and perirhinal regions).
